# Cryo-EM structures of human SID-1 transmembrane family proteins and implications for their low-pH-dependent RNA transport activity

**DOI:** 10.1101/2023.09.25.559159

**Authors:** Le Zheng, Tingting Yang, Hangtian Guo, Chen Qi, Yuchi Lu, Haonan Xiao, Yan Gao, Yue Liu, Yixuan Yang, Mengru Zhou, Henry C. Nguyen, Yun Zhu, Fei Sun, Chen-yu Zhang, Xiaoyun Ji

## Abstract

Human SIDT1 and SIDT2 are closely related members of the systemic RNA interference (RNAi)-defective (SID-1) transmembrane family. Both mediate RNA internalization and intracellular transport and are involved in various biological processes. However, the molecular basis of RNA uptake, especially for exogenous small RNAs, remains elusive. Here, we present the cryo-electron microscopy (cryo-EM) structures of human SIDT1 and SIDT2. Both structures reveal the overall architecture of a dimeric arrangement contributed by the β-strand-rich extracellular domain (ECD) and the transmembrane domain (TMD) with 11 passes, highlighting the remarkable structural congruence. The *in situ* assays confirm that SIDT1 and SIDT2 exist as dimers or higher-order oligomers. We demonstrate that for both SIDT1 and SIDT2, the ECD binds small RNAs, such as dietary plant-derived miRNA, only under acidic conditions. In addition, RNA binding under low pH can trigger higher-order assembly of the ECD dimer, suggesting the potential importance of oligomerization during RNA uptake. Our results illustrate the molecular features of the conserved SID-1 family proteins to elucidate the mechanism of the low pH-dependent activation of RNA uptake mediated by SIDT1 and SIDT2. This study provides a promising understanding of the molecular basis of the nucleic acid delivery platform, which may potentially open new avenues for the design and optimization of RNA-based therapies.

## Introduction

In the nematode *C. elegans*, the systemic RNA interference defective protein 1 (SID-1) plays a crucial role in systemic RNAi by facilitating the transport of exogenous double-stranded RNA (dsRNA) into the cytoplasm^1–5^. The human orthologs SIDT1 and SIDT2 belong to the mammalian SID-1 transmembrane family and share similarities with *C. elegans* SID-1, particularly in their nucleic acid transport ability^6^. Over the past decade, the involvement of SIDT1 and SIDT2 has been reported in a variety of biological processes, including glucose^7–9^ and lipid metabolism^10, 11^, innate immunity^12^, and tumorigenesis^13, 14^. However, their precise roles in these diverse processes have not been elucidated. Notably, SIDT1 and SIDT2 have been implicated in the regulation of these cellular functions through their involvement in RNA uptake and intracellular trafficking^15–17^.

In addition to endogenous microRNA (miRNA) playing important roles in cells, recent discoveries have shed light on the fascinating and trending topic of exogenous miRNAs, which regulate species from different kingdoms. SIDT1 has been shown to be predominantly localized to the plasma membrane and may function as an RNA transporter, particularly in facilitating RNA uptake^18^. For instance, overexpression of SIDT1 in pancreatic ductal adenocarcinoma cells enhances the uptake of synthetic small interfering RNA (siRNA) from the extracellular environment and strengthens siRNA-soaking-induced RNAi^6^. The discovery of natural cross-kingdom RNAi suggests that small RNAs act as signaling molecules that enable communication between organisms of different species and phylogenetic kingdoms^19–21^. Recently, dietary small RNAs, such as miRNAs have been shown to be absorbed through the mammalian digestive system and regulate host gene expression^22–25^. Our previous studies have discovered that SIDT1-deficient (*Sidt1^−/−^*) mouse models significantly impair the absorption of plant-derived miRNA in the stomach, and the acidic environment can enhance the SIDT1-mediated absorption of plant-derived miRNAs through unknown mechanisms^18^. Notably, the uptake of mature miRNAs by SIDT1 was significantly enhanced under acidic culture conditions, while the effect on double-stranded miRNA mimics was not significant^18^. Dietary miRNAs absorbed via SIDT1 could enter extracellular vesicles, such as exosomes, and affect target organs in the host^18^. Interestingly, a SIDT1 polymorphism carrying a V78M mutation affected the uptake of therapeutic miRNA in a clinical trial^26^.

Similarly, SIDT2 enables the uptake of naked single-stranded oligonucleotides into cells^27^. SIDT2 is localized to the lysosomal membrane and plays a role in activating the innate immune response by facilitating the transport of internalized dsRNA across the lysosome^12^. This then enables engagement with cytoplasmic RIG-I-like receptors (RLRs) and subsequently induction of type I interferon (IFN) production^12^. Comparable results have also been reported for SIDT1^15, 28^. SIDT2 plays a dual role in RNautophagy or DNautophagy by mediating the translocation of both RNA and DNA to lysosomes for degradation^13, 17, 29–31^. Recent studies have also uncovered an additional function of SIDT2 as a ceramidase to hydrolyze lipids, which may be involved in RNA uptake processing^32^.

Similar to SID-1, both SIDT1 and SIDT2 are predicted to have an extensive N-terminal ECD, a TMD with 11-transmembrane helices (TMs), and a flexible intracellular domain (ICD) loop of approximately 100 amino acids between the first two transmembrane helices^4, 33^. Loss of function studies focusing on RNA transport of SID-1 have highlighted the essential role of both the ECD and TMD in SID-1-mediated RNA transport function, with the ECD appearing to play a critical role in RNA binding and subsequent transport processes^33^. Although SIDT1- and SIDT2-dependent RNA uptake has been observed in various *in vitro* and *in vivo* systems^12, 18, 29, 30, 34^, the exact mechanisms underlying this uptake, particularly substrate recognition by SIDT1 and SIDT2, remain elusive.

In this study, we present the cryo-EM structures of human SIDT1 and SIDT2. Comparisons of these two structures reveal their overall architecture consisting of a dimeric conformation and highlight their structural congruence. *In situ* assays confirmed the existence of SIDT1 and SIDT2 as dimers or higher-order oligomers. Combined with biochemical assays, our results demonstrate the ability of SIDT1 and SIDT2 to interact with small RNAs, such as miRNA, in a pH-dependent manner under low pH conditions. Additionally, the binding of small RNAs under acidic conditions further triggers the ECD oligomerization. Taken together, these findings elucidate the molecular features of mammalian SID-1 transmembrane family proteins and provide compelling evidence for the involvement of SIDT1 in the uptake of exogenous small RNAs. Our work has implications for understanding the mechanism of low pH-dependent RNA uptake by SIDT1 and SIDT2.

## Results

### Purification and cryo-EM structure determination of human SIDT1 and SIDT2

The expression and purification of homogeneous membrane protein samples have been a major bottleneck in the structure determination of human SID transmembrane family proteins. To improve both expression yield and biochemical stability, we screened various constructs for SIDT1 and SIDT2. Eventually, we expressed SIDT1 and SIDT2 in *Spodoptera frugiperda* 9 (*Sf*9) cells, yielding protein samples with optimal solution behavior. After affinity purification, the presence of SIDT1 and SIDT2 was confirmed by Coomassie blue staining of SDS-polyacrylamide gel electrophoresis (SDS-PAGE) analysis. Subsequently, the peak fractions of the *Sf*9-expressed protein with high purity and homogeneity obtained from gel filtration chromatography were pooled and concentrated for cryo-EM analysis (Supplementary Fig. 1a-d).

The cryo-EM structures of SIDT1 and SIDT2 at pH 7.5 were determined at overall resolutions of 3.33 Å and 3.17 Å, respectively (Supplementary Figs. 2 and 3, and Table 1). Detailed information on protein purification, sample preparation, cryo-EM data acquisition and processing, and model building are described in Methods, as well as Supplementary Figs. 2 to 4 and Table 1. These high-quality density maps were used to build structural models and facilitate in-depth structural analysis. The electron density map of SIDT1 exhibited remarkable main-chain connectivity and well-defined side-chain densities for most residues within the SIDT1 ECD (Supplementary Fig. 4a). Similarly, the cryo-EM map of the SIDT2 ECD exhibited high integrity and quality, facilitating an accurate alignment between the structural model and the electron density (Supplementary Fig. 4b). However, the densities corresponding to the TMs of both SIDT1 and SIDT2 were refined to a lower resolution, with TM6 of SIDT1 exhibiting quite poor densities that hindered accurate model building (Supplementary Fig. 4c, d). The overall RMSD between our cryo-EM structures of SIDT1 and SIDT2 was 0.92 Å, with the structural differences primarily located in the TMD region. These findings further emphasize the substantial similarity in protein characteristics between SIDT1 and SIDT2, highlighting a significant degree of homology between the two proteins.

**Table 1.**
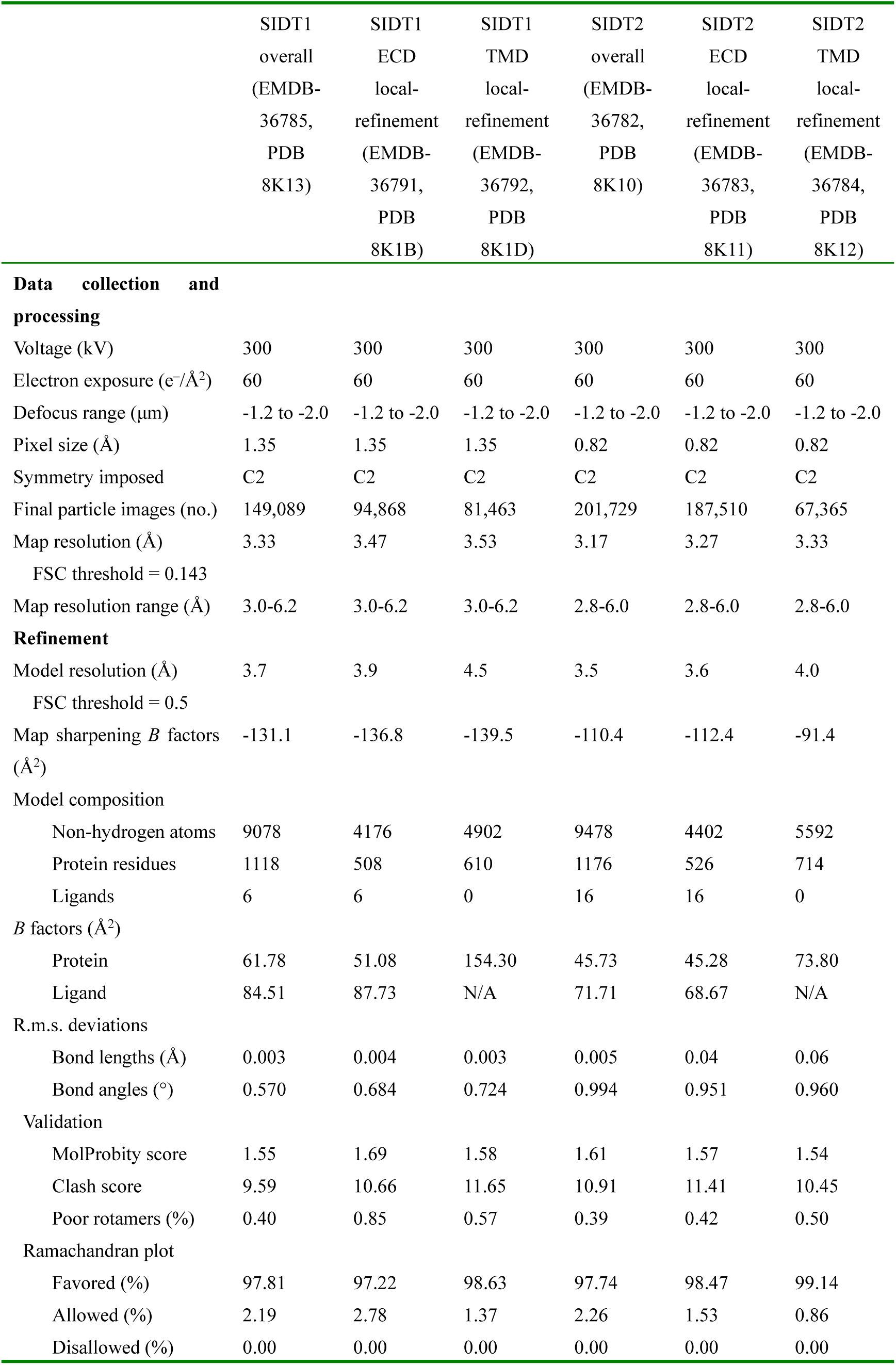
Cryo-EM data collection, refinement and validation statistics.

|  | SIDT1<br>overall<br>(EMDB-36785,<br>PDB<br>8K13) | SIDT1<br>ECD<br>local-<br>refinement<br>(EMDB-36791,<br>PDB<br>8K1B) | SIDT1<br>TMD<br>local-<br>refinement<br>(EMDB-36792,<br>PDB<br>8K1D) | SIDT2<br>overall<br>(EMDB-36782,<br>PDB<br>8K10) | SIDT2<br>ECD<br>local-<br>refinement<br>(EMDB-36783,<br>PDB<br>8K11) | SIDT2<br>TMD<br>local-<br>refinement<br>(EMDB-36784,<br>PDB<br>8K12) |
| --- | --- | --- | --- | --- | --- | --- |
| **Data collection and processing** |  |  |  |  |  |  |
| Voltage (kV) | 300 | 300 | 300 | 300 | 300 | 300 |
| Electron exposure (e<sup>-</sup>/Å<sup>2</sup>) | 60 | 60 | 60 | 60 | 60 | 60 |
| Defocus range (μm) | -1.2 to -2.0 | -1.2 to -2.0 | -1.2 to -2.0 | -1.2 to -2.0 | -1.2 to -2.0 | -1.2 to -2.0 |
| Pixel size (Å) | 1.35 | 1.35 | 1.35 | 0.82 | 0.82 | 0.82 |
| Symmetry imposed | C2 | C2 | C2 | C2 | C2 | C2 |
| Final particle images (no.) | 149,089 | 94,868 | 81,463 | 201,729 | 187,510 | 67,365 |
| Map resolution (Å) | 3.33 | 3.47 | 3.53 | 3.17 | 3.27 | 3.33 |
| FSC threshold = 0.143 |  |  |  |  |  |  |
| Map resolution range (Å) | 3.0-6.2 | 3.0-6.2 | 3.0-6.2 | 2.8-6.0 | 2.8-6.0 | 2.8-6.0 |
| **Refinement** |  |  |  |  |  |  |
| Model resolution (Å) | 3.7 | 3.9 | 4.5 | 3.5 | 3.6 | 4.0 |
| FSC threshold = 0.5 |  |  |  |  |  |  |
| Map sharpening *B* factors (Å<sup>2</sup>) | -131.1 | -136.8 | -139.5 | -110.4 | -112.4 | -91.4 |
| **Model composition** |  |  |  |  |  |  |
| Non-hydrogen atoms | 9078 | 4176 | 4902 | 9478 | 4402 | 5592 |
| Protein residues | 1118 | 508 | 610 | 1176 | 526 | 714 |
| Ligands | 6 | 6 | 0 | 16 | 16 | 0 |
| ***B* factors (Å<sup>2</sup>)** |  |  |  |  |  |  |
| Protein | 61.78 | 51.08 | 154.30 | 45.73 | 45.28 | 73.80 |
| Ligand | 84.51 | 87.73 | N/A | 71.71 | 68.67 | N/A |
| **R.m.s. deviations** |  |  |  |  |  |  |
| Bond lengths (Å) | 0.003 | 0.004 | 0.003 | 0.005 | 0.04 | 0.06 |
| Bond angles (°) | 0.570 | 0.684 | 0.724 | 0.994 | 0.951 | 0.960 |
| **Validation** |  |  |  |  |  |  |
| MolProbity score | 1.55 | 1.69 | 1.58 | 1.61 | 1.57 | 1.54 |
| Clash score | 9.59 | 10.66 | 11.65 | 10.91 | 11.41 | 10.45 |
| Poor rotamers (%) | 0.40 | 0.85 | 0.57 | 0.39 | 0.42 | 0.50 |
| **Ramachandran plot** |  |  |  |  |  |  |
| Favored (%) | 97.81 | 97.22 | 98.63 | 97.74 | 98.47 | 99.14 |
| Allowed (%) | 2.19 | 2.78 | 1.37 | 2.26 | 1.53 | 0.86 |
| Disallowed (%) | 0.00 | 0.00 | 0.00 | 0.00 | 0.00 | 0.00 |

### The overall structures of the human SIDT1 and SIDT2 homodimers

Previous studies have suggested that all members of the SID-1 transmembrane family are estimated to have 8-11 transmembrane segments^4, 33, 35^. Consistent with secondary structure predictions, SIDT1 and SIDT2 comprises a signal peptide (SP) at its N-terminal region, an ECD composed of two β-strand rich subdomains (ECD1 and ECD2), a large flexible ICD connecting TM1 and TM2, and a total of 11 transmembrane helices (Fig. 1a, b). Interestingly, SIDT1 forms a homo-dimeric structure involving its ECD and TMD that closely resembles the SIDT2 dimer structure, except for a slightly larger ICD in SIDT2 by approximately 10 amino acids (Fig. 1c, d). Both SIDT1 and SIDT2 dimeric proteins adopt highly symmetric forms with a two-fold rotational symmetry axis perpendicular to the membrane surface (Fig. 1c, d). To clarify their oligomeric states in solution, blue native gel assays were performed with purified full-length SIDT1 and SIDT2. The results indicated that both SIDT1 and SIDT2 exhibit a dimeric conformation of approximately 240 kDa. The molecular weights are influenced by extensive *N*-glycosylation modifications (Supplementary Fig. 1e). These results corroborate our structures of SIDT1 and SIDT2 homodimers.

**Fig. 1.**
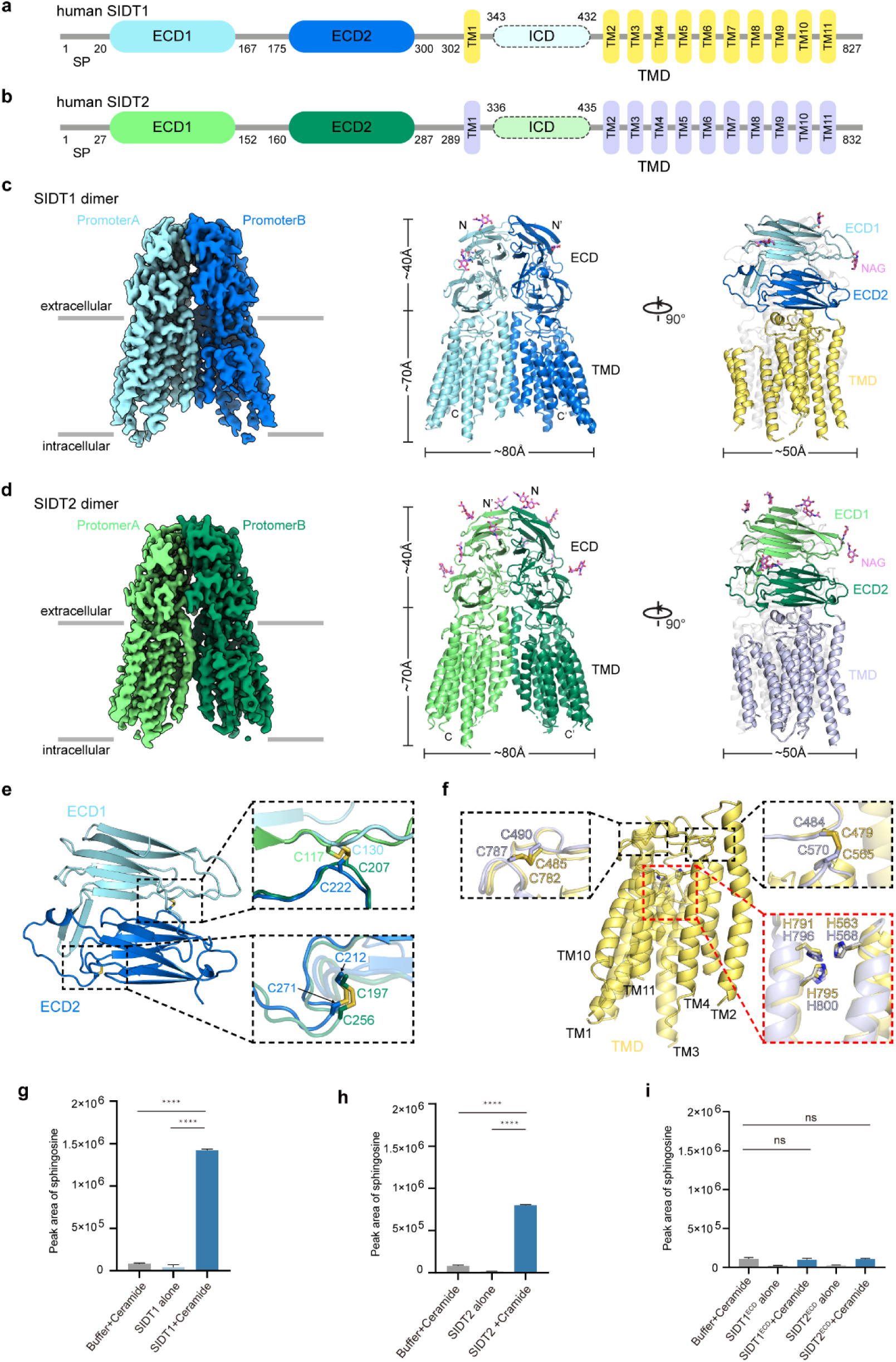
Overall structures of human SIDT1 and SIDT2, and LC-MS/MS analysis of their ceramidase activity. **a, b** Overview diagrams of the domain arrangement of the SIDT1 (**a**) and SIDT2 (**b**). SP, signal peptide; ECD, extracellular domain; ICD, intracellular domain; TMD, transmembrane domain. **c, d** Overall cryo-EM maps and structure models of the SIDT1 (**c**) and SIDT2 (**d**). Both SIDT1 and SIDT2 are homodimers. Protomers A and B in SIDT1 are colored light blue and deep blue, while in SIDT2 are colored light green and deep green, respectively. **e** A close-up view of the SIDT1 ECD. The SIDT1 ECD contains two subdomains, ECD1 and ECD2. Two pairs of conserved disulfide bonds that enhance the structural stability of the ECD are shown as sticks, along with the corresponding disulfide bonds in SIDT2. Both SIDT1 and SIDT2 models are colored according to the same scheme in (**c**) and (**d**). **f** A close-up view of the SIDT1 TMD. Two pairs of disulfide bonds that connected the loop 2-3 to loop 4-5 and loop 10-11 are shown as sticks, along with the corresponding disulfide bonds in SIDT2. The putative Zn^2+^-binding site is also conserved in SIDT1 and SIDT2 and the corresponding three histidine residues are shown as sticks. Both SIDT1 and SIDT2 models are colored according to the same scheme in (**c**) and (**d**). **g-i** Quantification of the ceramidase activity of SIDT1 (**g**), SIDT2 (**h**) and SIDT1/2ECD (**i**) was performed using d-erythrosphingosine (d18:1) as a standard through high-resolution LC-MS/MS analysis.

Bioinformatics analysis has identified eight predicted *N*-glycosylation sites in SIDT1, five of which are located on the ECD. Notably, the glycosylation sites at N67, N83, and N136 have been resolved in the EM map (Fig. 1c), while the precise determination of the remaining glycosylation sites was limited by the quality of the electron density. ECD1 (residues 20-167) adopts a 10-segment β-strands structure, including 4 glycosylation sites. ECD2 (residues 175 to 300) consists of 8 β-strands and 1 glycosylation site. ECD1 and ECD2 of SIDT1 are connected by a flexible loop, and their interaction is further reinforced by a disulfide bond formed between C130 of ECD1 and C222 of ECD2, which contributes to the overall stability of SIDT1 (Fig. 1e and Supplementary Fig. 4e). Additionally, another pair of disulfide bonds between C212 and C271 on ECD2 further contributes to its structural integrity (Fig. 1e and Supplementary Fig. 4e). SIDT2 also features ten predicted *N*-glycosylation sites, with all six glycosylation sites (N27, N54, N60, N123, N141, and N165) on its ECD well resolved in the EM map (Fig. 1d). In addition, two pairs of disulfide bonds are also present at the corresponding positions in the SIDT2 ECD (Fig. 1e and Supplementary Fig. 4f).

The SIDT1 TMD consists of 11 transmembrane helices, each playing an important role in the formation of the stable homodimer. Similar to the ECD, the TMD also has two pairs of disulfide bonds (Fig. 1f and Supplementary Fig. 4g). One pair, formed by C479 and C565, connects the loop between TM2 and TM3 (loop 2-3) to the loop between TM4 and TM5 (loop 4-5). The other pair, formed by C485 and C782, connects loop 2-3 to the loop between TM10 and TM11 (loop 10-11). Importantly, these two pairs of conserved disulfide bonds are also present at corresponding positions within the SIDT2 TMD, highlighting a common structural feature in the SID-1 transmembrane family (Fig. 1f and Supplementary Fig. 4h). Indeed, the conservation of disulfide bonds extends to the entire SID-1 transmembrane family, including SIDT1 and SIDT2 (Supplementary Fig. 5). This highlights the significance of these disulfide bonds in maintaining structural integrity and functional properties within the family.

Notably, a recent structural study on SIDT2 emphasized the existence of three histidine residues (H568, H796 and H800) within its TMD, which was responsible for lipid hydrolytic activity^32^. These histidine residues, together with S564 in TM4 and D579 in TM5, potentially form a zinc-binding site, as supported by the EM map displaying a potential density for a zinc ion^32^. Although the presence of zinc ions could not be confirmed in our EM map of SIDT1 due to resolution limitations and lack of relevant functional experiments, we did observe three histidine residues (H563 in TM4 and H791 and H795 in TM11) at corresponding positions in SIDT1. The findings in SIDT2 (Fig. 1f) provide strong support for our observations. Furthermore, our biochemical assays successfully confirmed the ceramidase activity of both SIDT1 and SIDT2 (Fig. 1g-I and Supplementary Fig. 6). These structural similarities serve to emphasize the homology and functional relevance between SIDT1 and SIDT2.

### Comparison of homodimer formation of human SIDT1 and SIDT2

Both the ECD and the TMD of SIDT1 are essential for its homodimer interactions. The dimer interface on the ECD can be divided into two regions (Fig. 2a). The first region is formed by the ECD1 subdomains from two protomers, mainly through extensive hydrophobic interactions (Fig. 2b). Additional hydrogen bond interactions also contribute to the stability of the dimeric structure, including the conserved hydrogen bond formed between S105 residues in both protomers. This conserved hydrogen bond is also observed in SIDT2, where the corresponding amino acid is S92 (Fig. 2b). Another dimer interface region mediated by the ECD involves the interaction of a long loop on ECD2 that connects two β-strand sheets (Fig. 2c). Notably, a possible hydrogen bond interaction exists between H229 in one protomer and N230 in the other protomer (Fig. 2c), which closely resembles the interaction observed in SIDT2, although the histidine is replaced by asparagine (N214 in SIDT2). A recent structural study of SIDT2 also reported that Y210 and F218 from one protomer can form cation-π interactions with R100 from the opposing protomer, and D204 from one protomer forms ionic interactions with both R100 and K106 from the other protomer^32^. We did not observe such an interaction in the structure of SIDT1, where the Q113 in SIDT1, corresponding to R100 in SIDT2, has a shorter side chain, making it unlikely to interact with the corresponding Y225 and F233. However, Q113 may maintain a hydrogen bond interaction with N219 from the other protomer instead of the observed ionic bond interaction observed between R100 and D204 in SIDT2 (Fig. 2c). This may result in the dimer interface of SIDT1 being less enriched in interactions compared to SIDT2.

**Fig. 2.**
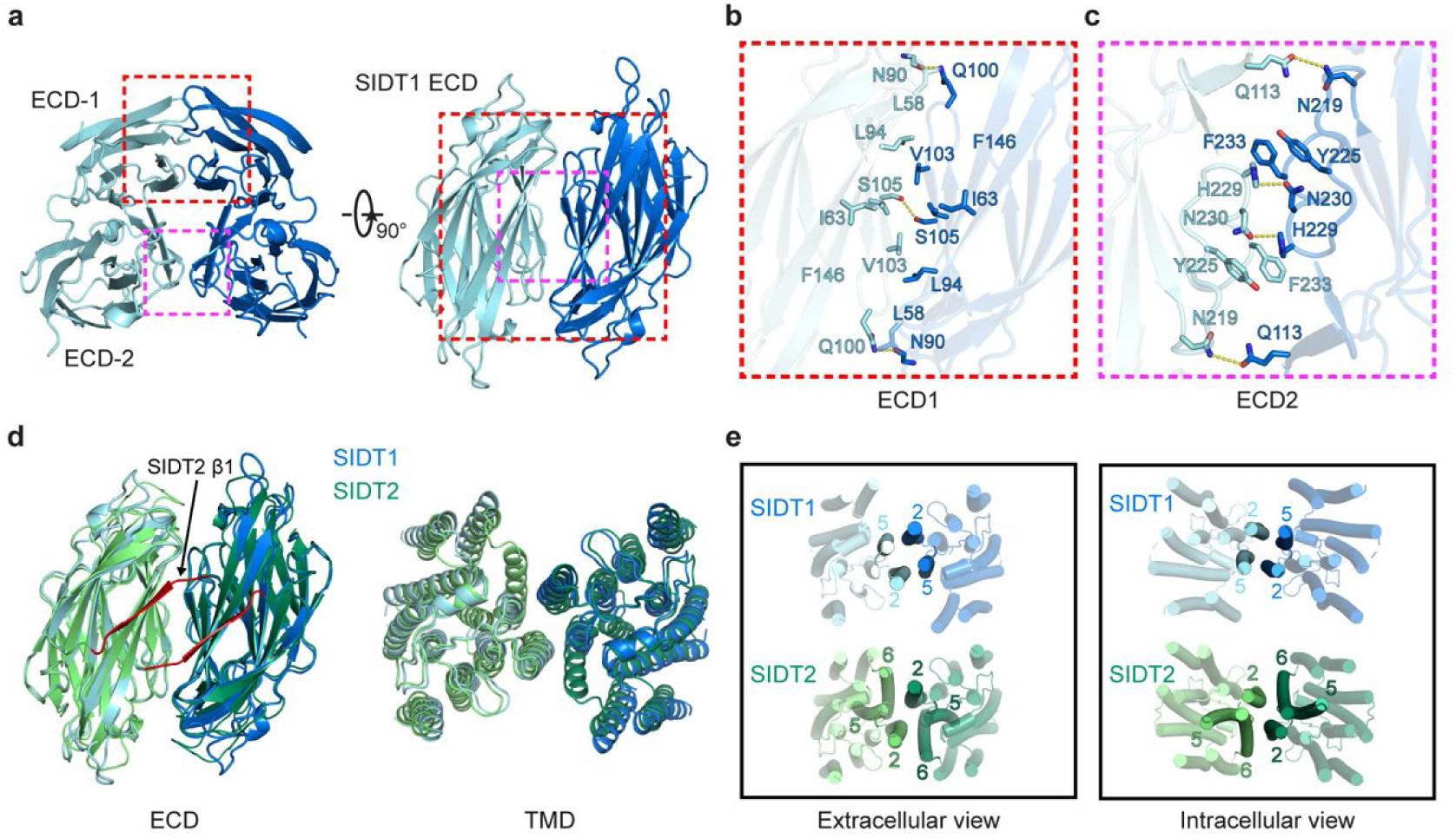
Detailed interactions in the dimer interface of SIDT1. **a** The dimer interface on SIDT1 ECD contains two main regions (boxed with red and magenta dashed lines, respectively). **b, c** The detailed dimeric interactions in ECD1 (**b**) and ECD2 (**c**) between two SIDT1 protomers. The potential hydrophobic interacting residues are shown as sticks. Potential hydrogen bonds are represented as yellow dashed lines. **d** Comparison of the dimer interface between SIDT1 and SIDT2, including the ECD (left) and the TMD (right). The first β-strand (β1) at the N-terminal of SIDT2 is colored red. **e** The dimer interface in the SIDT1 TMD is mainly mediated by TM2, which interacts with TM5 of the other protomer, while dimeric interactions in the SIDT2 TMD primarily involve TM2 and TM6.

The dimerization interaction involves a buried surface area of ∼1700 Å^2^ from each SIDT2 protomer, which is consistent with the recently published SIDT2 structure (PDB ID: 7Y68)^32^. In contrast, the dimer interface of the ECD interaction measured in our SIDT1 model is ∼1300 Å^2^ in each protomer, suggesting that while the structures of SIDT1 and SIDT2 are highly homologous and share similar structural features, there are distinct differences in the proximity of the homodimers formed by them, probably due to their sequence variations. SIDT2 contains an additional β-strand (residues 27 to 33) that participates in interactions at the dimer interface (Fig. 2d). In contrast, the corresponding region in SIDT1 is unstructured, indicating a high level of flexibility.

Compared to the ECD, the overall arrangement of the TMDs in both SIDT1 and SIDT2 is similar, but different TMs contribute to dimerization (Fig. 2d). In SIDT1, the dimer interface is primarily mediated by TM2, which interacts with TM5 of the other protomer (Fig. 2e). Despite the absence of precise localization of TM6 in our structure model, it is unlikely that TM6 plays a significant role in the dimer interface interaction of SIDT1, as inferred from the relative positions of TM5 and TM7. These interactions are largely driven by extensive hydrophobic interactions. In SIDT2, however, the interactions primarily involve TM2 and TM6, which represents a key distinction between the TMDs of SIDT1 and SIDT2 (Fig. 2e). Overall, while the ECD structure exhibits a high degree of consistency, the individual helices of the TMD in SID-1 transmembrane family members show a degree of flexibility, highlighting the dynamic nature of these proteins in performing specific functions at the membrane.

### SIDT1 exhibits oligomeric states *in situ* and *in vitro*

Previous studies have shown that co-expression of a missense mutant SID-1 protein interferes with wild-type SID-1 function in a dose-dependent manner, suggesting that SID-1 functions as a oligomer^2^. Here, our cryo-EM structures of SIDT1 and SIDT2 reveal a homodimeric arrangement in micelles. Therefore, we investigated the *in situ* oligomeric states of SIDT1 and SIDT2 in living cells using a bimolecular fluorescence complementation (BiFC) assay. We co-expressed SIDT1 or SIDT2 tagged with two complementary non-fluorescent fragments of green fluorescent protein (GFP) in living cells and monitored the proper assembly of the complete GFP (Fig. 3a). The BiFC results showed robust green fluorescence signal in living cells, indicating that both SIDT1 and SIDT2 have the ability to form dimers or oligomers (Fig. 3b, c). We also tested the TMD truncations of SIDT1 and SIDT2 in the BiFC assay and found GFP fluorescence intensity comparable to those observed for full-length proteins (Fig. 3b, c), suggesting that TMD can dimerize or multimerize even in the absence of the ECD.

**Fig. 3.**
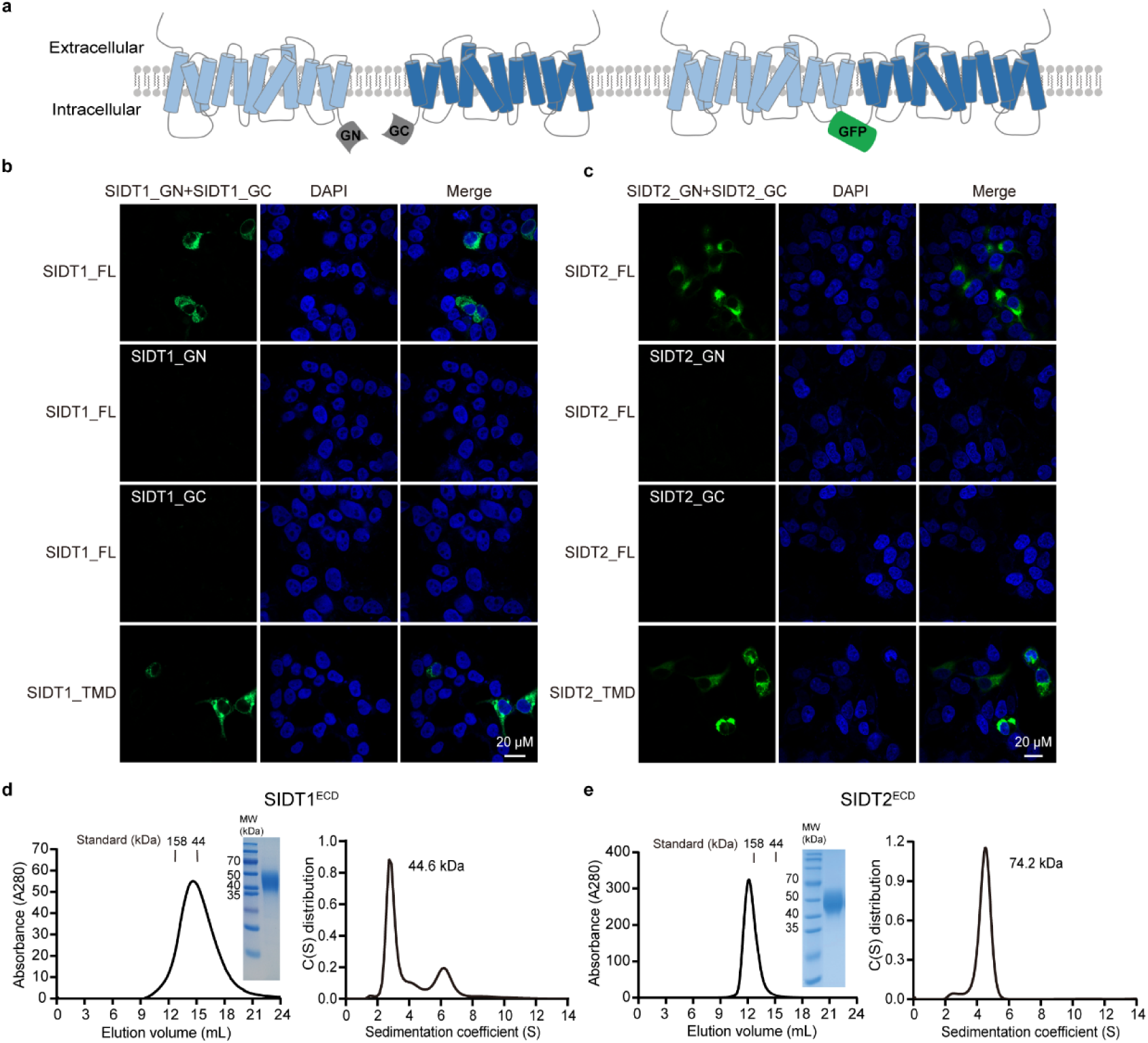
SIDT1 exhibits oligomeric states *in situ* and *in vitro*. **a** Schematic diagram representing the principle of the BiFC assay. **b, c** High-resolution confocal images of HEK293T cells expressing SIDT1 or SIDT2 (FL: full-length, top; TMD, bottom) fused to the N-terminal and C-terminal fragments of GFP fluorescent protein. Scale bar, 20 µm. **d, e** Characterization of the purified SIDT1^ECD^ (**d**) and SIDT2^ECD^ (**e**). The left panel displays the SEC of purified SIDT1^ECD^ using a Superdex 200 increase 10/300 GL column and the SDS-PAGE analysis of peak fractions of interest. The right panel shows SV-AUC analysis of the molecular weight of SIDT1^ECD^ (**d**) and SIDT2^ECD^ (**e**) in solution. These SV-AUC experiments were performed twice with equivalent results. Values in the panel represent mean ± s.d.

In addition, we investigated the role of the ECD in the oligomerization states of the proteins. We expressed and purified ECD truncations of both SIDT1 (SIDT1^ECD^) and SIDT2 (SIDT2^ECD^) in FreeStyle293-F cells and analyzed the homogeneous fractions by size exclusion chromatography (SEC). The elution volume for SIDT1^ECD^ was 14.6 mL, while that for SIDT2^ECD^ was 12.5 mL (Fig. 3d, e), suggesting that their oligomerization status may be different. For independent verification, we measured the sedimentation rate of the proteins by sedimentation velocity analytical ultracentrifugation (SV-AUC). SIDT1^ECD^ sedimented with a coefficient of 2.8 S and an approximate molecular weight of 44.6 kDa, indicating a monomer. Conversely, SIDT2^ECD^ was detected as a dimer with a sedimentation coefficient of 4.8 S and an estimated molecular weight of 74.2 kDa (Fig. 3d, e). The distinct oligomerization status of ECD is likely due to their varied amino acid composition and the subsequent interactions between the dimer interface with different buried areas, which is consistent with what we have observed in the structures, despite a highly conserved fold.

Taken together, SIDT1 and SIDT2 exist as dimers or oligomers, and the TMD plays a critical role in maintaining the dimeric assemblies.

### ECD of SIDT1 and SIDT2 bind small RNAs in a pH-dependent manner

The ECD of SID-1, SIDT1 and SIDT2 have been previously studied for their RNA binding capacity^33^. However, it has been reported that both SIDT1 and SIDT2 ECDs bind only long dsRNA, but not dsRNA shorter than 100 bp^16^. This conflicts with the function of the SID-1 transmembrane family which mediate the uptake of exogenous small RNAs. To examine this, we performed an electrophoretic mobility shift assay (EMSA) to examine the RNA binding ability of SIDT1^ECD^ using a synthetic miR168a, a 21 nt plant miRNA, previously demonstrated to be internalized by SIDT1^23^. The miR168a duplex was also used in the assay, termed dsmiR168a. At pH 8.0, SIDT1^ECD^ showed no detectable binding to either single-stranded (ss) or dsmiR168a (Fig. 4a). Interestingly, when the pH was lowered to 5.5, SIDT1^ECD^ showed clear binding to both forms of miR168a (Fig. 4a). Considering the dimerization status of full-length SIDT1 compared to monomeric SIDT1^ECD^, we engineered a fusion protein of SIDT1^ECD^ with immunoglobulin fragment crystallizable region (Fc; SIDT1^ECD-Fc^) to enhance the dimerization of SIDT1^ECD^ (referred to as SIDT1^ECD-Dimer^). SIDT1^ECD-Dimer^ was stabilized by Fc dimerization, and confirmed by SV-AUC (Supplementary Fig. 7a, b). The EMSA results showed essentially the same RNA binding feature for SIDT1^ECD-Dimer^ as for SIDT1^ECD^ (Fig. 4b).

**Fig. 4.**
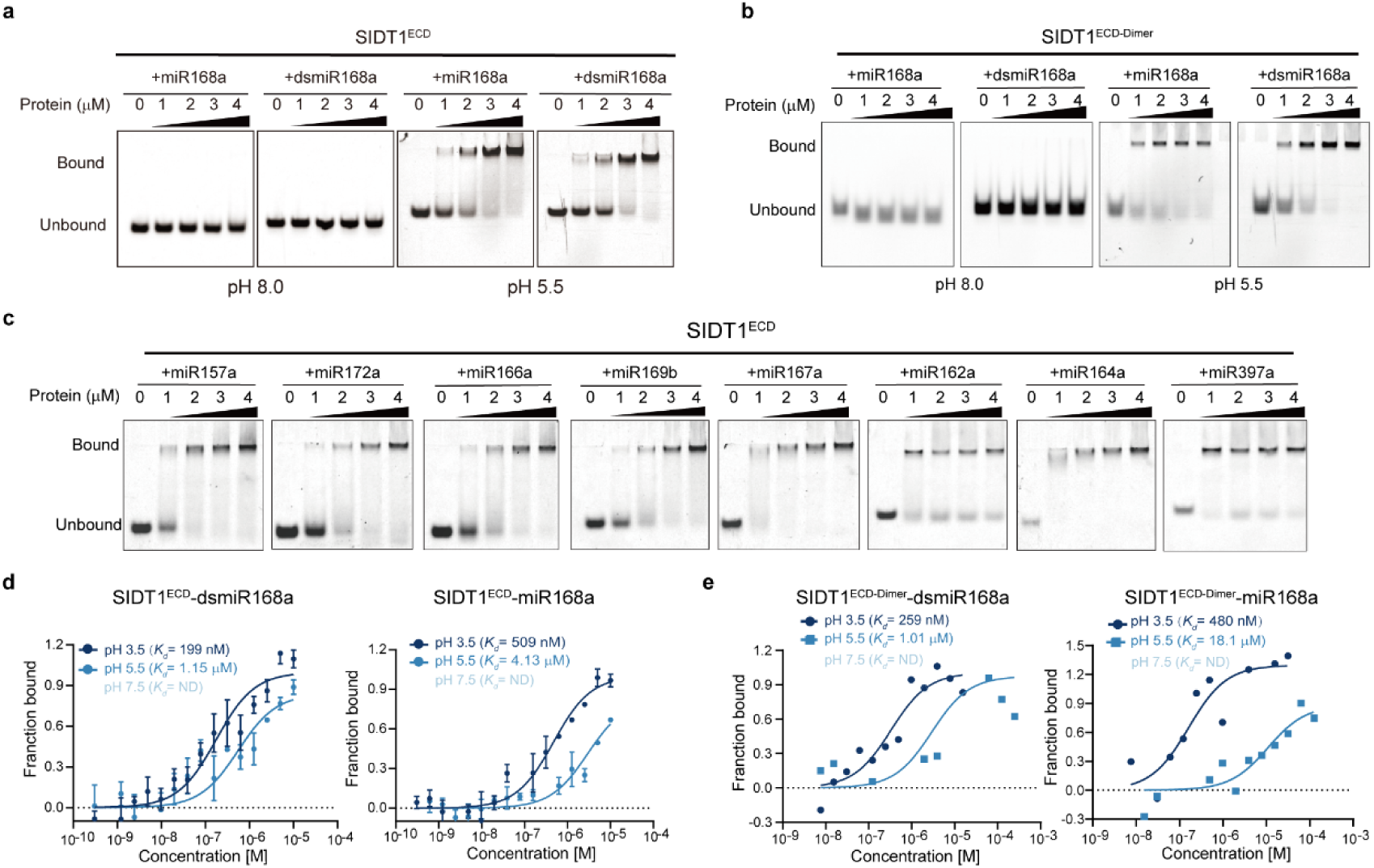
SIDT1^ECD^ binds to small RNAs under acidic conditions and exhibits a pH-dependent manner. **a, b** EMSA results of SIDT1^ECD^ (**a**) or SIDT1^ECD-Dimer^ (**b**) under pH 5.5 and pH 8.0 conditions. The final protein concentrations in lanes 1-5 are 0, 1, 2, 3, and 4 μM, respectively, and the final concentrations of the 5’-FAM-labeled miRNAs are 2.5 μM in each lane. Bound, protein-miRNA complexes; Unbound, free miRNAs. **c** SIDT1^ECD^ binds to various plant-derived miRNAs encoding different sequences at pH 5.5. Plant-derived miRNA sequences are listed in Supplementary Table 1. **d, e** MST analysis measuring binding affinities of SIDT1^ECD^ (**d**) and SIDT1^ECD-Dimer^ (**e**) with small RNAs across different pH conditions. The calculated dissociation constant (*K_d_*) value represents the affinity between SIDT1^ECD^ or SIDT1^ECD-Dimer^ and miRNAs. All MST data results are the means of three independent experiments.

Next, we examined the binding of SIDT1^ECD^ and SIDT1^ECD-Dimer^ to a panel of plant-derived miRNAs prevalent in the serum of healthy Chinese individuals^23, 36^ at pH 5.5. The EMSA results showed that all the miRNAs tested could be bound to SIDT1 under acidic conditions (Fig. 4c and Supplementary Fig. 7c). We also synthesized corresponding DNA sequences and examined the interactions at pH 5.5. The EMSA results demonstrated that the ECD could also bind to both single-stranded DNA (ssDNA) and double-stranded DNA (dsDNA) (Supplementary Fig. 7d), indicating a lack of nucleic acid type specificity. The same experimental results were also confirmed in the case of SIDT2 (Supplementary Fig. 7e). Our results revealed that ECD, whether as a monomer or a dimer, has a clear ability to bind small nucleic acid *in vitro* and the binding is low-pH-dependent.

Next, we quantified the RNA binding affinity of ECD to miR168a and dsmiR168a under various pH conditions (pH 7.5, 5.5, and 3.5) using Microscale Thermophoresis (MST). MST results revealed that at pH 3.5, the dissociation constant (*K_d_*) of SIDT1^ECD^ for miR168a is 509 nM, while the *K_d_* for dsmiR168a is 199 nM. At pH 5.5 and with equal nucleic acid saturation, the detected *K_d_* between SIDT1^ECD^ and dsmiR168a was significantly increased to 1.15 μM. Furthermore, at pH 7.5, the interactions were too weak to establish a discernible binding curve (Fig. 4d). Similarly, MST analysis demonstrated that SIDT1^ECD-Dimer^ exhibits high affinity binding to both dsmiR168a and miR168a at pH 3.5, with *K_d_* of 259 nM and 480 nM, respectively. However, the binding affinity decreased significantly when the pH was increased to 5.5, resulting in *K_d_* of 1.01 μM and 18.1 μM for dsmiR168a and miR168a, respectively. In a neutral condition (pH 7.5), we were only able to detect extremely weak interactions to a limited extent (Fig. 4e), confirming the observations from our EMSA experiments (Fig. 4a, b).

When protein stability was measured by differential scanning fluorometry (DSF), the ECDs of SIDT1 and SIDT2 both exhibited enhanced thermostability under acidic conditions compared to neutral conditions (Supplementary Fig. 7f), highlighting the potential low pH-dependent functions of SIDT1 and SIDT2 in their physiologically acidic working environment.

Furthermore, we investigated the surface electrostatic potential of the ECD dimers of SIDT1 and SIDT2 structures at different pH values (Supplementary Fig. 8). Our analysis revealed the presence of positively charged regions on the β-strand surfaces of both SIDT1 and SIDT2 ECDs at pH 7.5. Interestingly, these positively charged regions exhibited an increased charge density and expanded distribution at pH 5.5, and nearly all regions of the ECDs in both SIDT1 and SIDT2 displayed a positive charge at pH 3.5. This pH-dependent variation in surface charge properties supports our EMSA data on the pH-dependent nucleic acid binding ability of SIDT1 and SIDT2. Taken together, our biochemical results combined with structural analysis revealed that the ECDs of SIDT1 and SIDT2 efficiently bind to small RNAs under acidic conditions.

### Acidic pH promotes RNA-mediated SIDT1 oligomerization

To further investigate the molecular impact of the pH-dependent interactions between ECD and RNAs, we investigated the formation and the biochemical properties of the RNA-ECD complexes. First, we subjected SIDT1^ECD-Dimer^ or SIDT1^ECD^ to analytical SEC and SV-AUC analysis under pH 3.5, 5.5, and 7.5, respectively. No significant differences in elution volume or sedimentation coefficient were observed at different pH values, indicating that acidic stimulation alone does not trigger higher-order oligomerization of SIDT1^ECD-Dimer^ and SIDT1^ECD^ (Fig. 5a, b and Supplementary Fig. 9a, b). Next, we incubated SIDT1^ECD-Dimer^ with dsmiR168a at pH 5.5 and obtained a stable complex during SEC analysis with an elution volume shifted forward by approximately 0.9 mL compared to SIDT1^ECD-Dimer^ alone (Fig. 5c), indicating a notable increase in molecular weight. An even more remarkable elution volume shift of 2.0 mL was observed when SIDT1^ECD^ was mixed with dsmiR168a (Fig. 5d), suggesting a possible formation of SIDT1^ECD^ tetramer. The SV-AUC analysis of SIDT1^ECD-Dimer^ bound to dsmiR168a at pH 5.5 yielded an average sedimentation coefficient of ∼9.5 S, indicating the formation of assemblies larger than dimers (Fig. 5e). SIDT1^ECD^ bound to dsmiR168a sedimented as a tetramer (∼150 kDa) with a substantially increased sedimentation coefficient of ∼4 S compared to the monomeric SIDT1^ECD^ (Fig. 5f). The SIDT1^ECD-Dimer^-dsmiR168a and SIDT1^ECD^-dsmiR168a complex formed at pH 5.5 was further cross-linked using glutaraldehyde and analyzed by SEC (Fig. 5g, h). SEC experiments confirmed that cross-linking of distinct higher-order oligomers of SIDT1ECD-Dimer and SIDT1ECD had occurred in the presence of RNA, as evidenced by a shift in the apparent molecular weight compared to the ECD in the absence of RNA (Fig. 5g, h). The cross-linked SIDT1^ECD^-dsmiR168a complex also revealed the formation of oligomers with RNA present (Supplementary Fig. 9c). The same experimental results were also corroborated in the case of SIDT2^ECD^ (Supplementary Fig. 9d). In conclusion, these results highlight the role of RNA in promoting ECD oligomerization under acidic conditions. It implies that SIDT1 and SIDT2 could form higher-order oligomers during low pH-dependent RNA uptake.

**Fig. 5.**
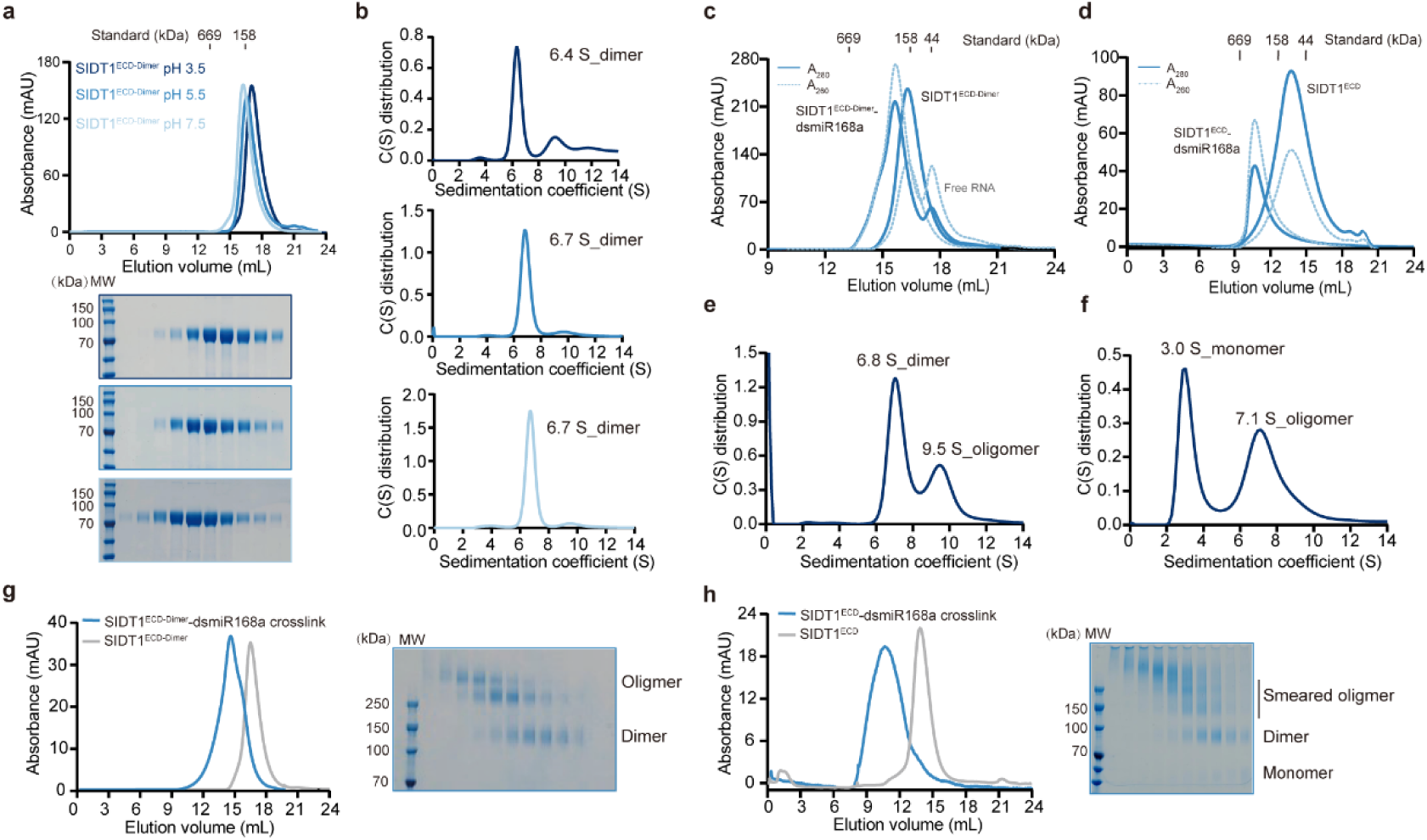
Acidic pH promotes RNA-mediated SIDT1 oligomerization. **a** SEC and SDS-PAGE analysis of SIDT1^ECD-Dimer^ at pH 3.5 (deep blue), pH 5.5 (blue), and pH 7.5 (light blue), respectively. **b** SV-AUC analysis of the sedimentation coefficient corresponding to SEC results. SV-AUC experiments were performed twice with equivalent results. Values in the panel represent mean ± s.d. **c, d** SEC analysis of SIDT1^ECD-Dimer^ (**c**) or SIDT1^ECD^ (**d**) forming complexes with dsmiR168a, respectively. Experiments were performed under pH 5.5 conditions. Blue solid line: A_280_; Light blue dashed line: A_260_. **e, f** SV-AUC analysis of the sedimentation coefficient of SIDT1^ECD-Dimer^-dsmiaR168a (**d**) and SIDT1^ECD^-dsmiaR168a (**f**) in solution under acidic conditions. Experiments were conducted in duplicate with comparable results. Panel values represent mean ± s.d. **g, h** SEC profiles of SIDT1^ECD-Dimer^ and SIDT1^ECD^ with dsmiR168a using non-reducing SDS-PAGE (**g**) or standard SDS-PAGE (**h**), respectively.

## Discussion

SIDT1 and SIDT2 are human orthologs of the *C. elegans* protein SID-1, which plays a pivotal role in systemic RNAi by transporting exogenous dsRNA into the cytoplasm^3–5^. SIDT1 and SIDT2 are involved in glucose and lipid metabolism, innate immunity, and tumorigenesis^8, 10, 12, 13, 28^. They have additional roles in cytokine regulation and inflammasome^14^ activation during the immune response which are linked to RNA uptake and intracellular trafficking. Despite the controversy surrounding the SIDT1-mediated absorption of exogenous small RNAs, it has initiated a new era of cross-kingdom regulation by plant-derived small RNAs^22^. Additionally, some studies have also indicated that SIDT1 possesses the ability to transport cholesterol^37^, while investigations into SIDT2 have uncovered lipid hydrolytic activity within its transmembrane domain^32, 38^. While further experimental evidence is needed to support these findings, they provide valuable insights into the multifaceted roles of SIDT1 and SIDT2 in cellular processes. Thus, the comprehensive understanding of the structures of SIDT1 and SIDT2 can serve as a solid foundation for future investigations into their functional mechanisms.

In this study, we have successfully determined the cryo-EM structures of human SIDT1 and SIDT2, providing valuable insights into their homodimer architectures. During the preparation of our manuscript, Qian et al. reported a human SIDT2 structure derived from FreeStyle 293-F cells^32^. In our study, we procured full-length SIDT1 and SIDT2 protein samples from both FreeStyle 293-F cells and insect *Sf9* cells. We ultimately chose sample data from the *Sf9* cells for the final structure determination of both SIDT1 and SIDT2. Our SIDT2 structure closely aligns with the recently published SIDT2 structure, indicating minimal disparities between proteins produced via two different expression systems.

The highly conserved features observed in the well-defined structures of SIDT1 and SIDT2, which share a 57% sequence identity^39^, provide valuable insights into their potential physiological roles. The ECD domain, featuring conserved β-sheet repeats, may display similar nucleic acid-binding interfaces and patterns, in line with our nucleic acid binding assays. While the ECD region in both SIDT1 and SIDT2 remains relatively rigid, SIDT2 features a larger and more tightly packed dimer interface (∼1700 Å^2^) compared to SIDT1 (∼1300 Å^2^), attributed to the unique amino acids at their interface. The V78M mutation (rs2271496), a clinically reported SIDT1 single nucleotide polymorphism (SNP), impacts exogenous small RNA uptake^26^ and resides on the β-strand within the ECD. This alteration may disrupt intricate interaction networks among neighboring amino acids, potentially reducing the ECD stability and impairing SIDT1-mediated nucleic acids binding and absorption. Additionally, the F169T and P186L mutations in SIDT1, and F154T and P171L mutations in the SIDT2^33^, are both situated in a flexible loop within the ECD and may influence the structural stability of the ECD, potentially affecting nucleic acid uptake by SIDT1. In contrast to the rigid ECD, the TMD displays increased flexibility. A comparison of experimentally determined SIDT1 and SIDT2 structures with their AlphaFold predicted counterparts reveal substantial variation in TMD arrangements, implying that TMD might undergo dynamic arrangement on the cell membrane^40^ (Supplementary Fig 10). This inherent flexibility may contribute to the nucleic acid uptake capabilities of the SID-1 transmembrane family.

Nematode studies have shed light on miRNA uptake in target cells, showing that *C. elegans* may utilize a distinctive SID-1-mediated dsRNA transfer pathway, essential for systemic RNAi and emphasizing the role of SID-1 in intercellular long dsRNA translocation^4, 5^. Previous research on SIDT1 and SIDT2 indicated that both ECDs bind 500- or 700-bp dsRNA, yet necessitate higher protein concentrations than SID-1 ECD^33^. Previous results suggest that at pH 7.0, SIDT1 ECD does not bind dsRNA shorter than 300 bp, while SIDT2 ECD exhibits no binding for dsRNA shorter than 100 bp^33^. Our study presents evidence that under acidic conditions (pH < 5.5), both SIDT1^ECD^ and SIDT2^ECD^ exhibit the capacity to indiscriminately bind small ds/ssRNAs rather than long dsRNA only in a pH-dependent manner, providing valuable supplementary information to prior research findings. Additionally, previous studies have shown that recombinant SID-1 ECD selectively binds dsRNA, not dsDNA, in a length-dependent manner at pH 7.0^33^. This conflicts with our findings of non-specific ECD-nucleic acid binding under acidic conditions, mainly due to differing pH values. It has been previously reported that SIDT1, localized on the apical membrane of gastric epithelial cells, works within the acidic gastric milieu (pH 2-4)^41^, wherein acid stimulation significantly enhances the uptake of exogenous small RNA^18^. In parallel, SIDT2, located on the lysosomal membrane (lysosomal lumen, pH 4.6-5.0)^42^, is involved in RNA or DNA internalization^29, 30^. Our results correspond with the acidic physiological conditions, indicating that under these conditions, both ECDs exhibit an affinity for small RNAs, offering a molecular basis for small RNAs uptake mediated by SIDT1 and SIDT2.

The protonation state of protein/RNA interface residues can play a crucial role in modulating their interactions, leading to pH-dependent events^43^ (Supplementary Fig. 7a, b). In our study, we speculate that the presence of histidine residues at the interface, known for their pKa of approximately 6.0, may contribute to the observed pH-dependent affinity between the ECD and RNA, particularly at pH 5.5 as observed in our biochemical assays (Supplementary Fig. 8c, d). Through structural analysis, we also identified co-localized, highly conserved charged residues (such as Arg, Lys, Asp, and Glu) on the ECD surface that are not involved in the dimer interface (Supplementary Fig. 5). These charged amino acids and histidine residues occupy the same surface location. Given the impact of low pH on histidine protonation, we propose that the ECD-RNA interaction primarily occurs at the back-to-dimer interface.

In addition, our research reveals that SIDT1^ECD^ binding to small RNAs under acidic conditions triggers higher-order oligomerization. However, the detection of SIDT1 oligomerization in our cryo-EM structure was hindered due to spatial constraints imposed by the presence of detergent micelles. Considering the TMD flexibility on the cell membrane, we speculate that SIDT1 may also exhibit oligomerization tendencies upon nucleic acid interaction (Supplementary Fig. 11). The oligomerization tendency holds the potential to play a crucial role in facilitating RNA uptake and even transport within the SID-1 transmembrane family, potentially involving additional interacting proteins or specific lipids. Further investigation is needed to explore these hypotheses. Despite the many obstacles and unresolved issues in the current research on SID-1 transmembrane family proteins, our investigation sheds additional light on the uncharted molecular feature of SID-1 transmembrane family proteins and highlights the potential influence of pH in controlling SIDT1-mediated RNA uptake. This study offers a deeper molecular foundation of this promising nucleic acid delivery platform, potentially impacting therapeutic applications.

## Methods

### Recombinant construct preparation

The full-length cDNAs for human SIDT1 and SIDT2 (Uniport: Q9NXL6 and Q8NBJ9, respectively) were codon-optimized and synthesized by GenScript Co., Ltd. To facilitate protein expression and structural analysis, we constructed internal truncated constructs of SIDT1 (SIDT1^ΔICD^, residues 366-408 deletion) and SIDT2 (SIDT2^ΔICD^, residues 366-410 deletion) in the pFastBac1 vector containing an N-terminal efficient signal peptide derived from the influenza virus hemagglutinin HA (MKANLLVLLCALAAADA) and a triple Flag tag sequence downstream of the coding region. Mutations were introduced using a two-step PCR-based strategy, followed by meticulous sequencing to confirm clone identities.

For the expression and purification of the ECD proteins of SIDT1 and SIDT2, we subcloned the DNA sequences corresponding to amino acids 23-310 of SIDT1 and 22-292 of SIDT2 into a customized pMlink vector. This vector contains an N-terminal melittin signal sequence and a C-terminal 6 × His-tag. To ensure stability, we utilized a fusion strategy by adding an Fc fragment to the C-terminus of the ECD to facilitate dimerization. We also introduced a 6 × His-tag and a short GS linker. The final construct was subcloned into the pMlink expression vector, and clone identities were verified by Sanger sequencing.

### Expression and purification of full-length SIDT1 and SIDT2

The full-length SIDT1 and SIDT2 were expressed using the Bac-to-Bac Baculovirus Expression System (Invitrogen). The *Sf9* cells were cultured in Sf-900 II SFM (Gibco, USA). Bacmid DNAs were generated and transfected into *Sf9* cells to produce and amplify baculovirus. A total of 6-8 liters of infected *Sf9* cells were harvested 60 hours post-infection and then lysed in lysis buffer containing 50 mM HEPES pH 7.5, 300 mM NaCl, and supplemented with cocktail inhibitor (1.04 mM AEBSF, 0.8 μM Aprotinin, 50 μM Bestatin, 15 μM E-64, 20 μM Leupeptin and 15 μM Pepstatin A, MCE, Cat# HY-K0010) and 1 mM PMSF. Cells were disrupted using high-pressure cell disruption at 800 bars. The cell debris was removed by low-speed centrifugation for 30 minutes and the supernatant was further subjected to high-speed centrifugation at 150,000 × g for 120 minutes using a Ti-45 rotor (Beckman). Then the membrane component was resuspended and homogenized using a glass dounce homogenizer for 20 cycles on ice in solubilization buffer containing 50 mM HEPES pH 7.5, 300 mM NaCl, 1 mM EDTA, 2% (*w/v*) *n*-dodecyl-β-D-maltoside (DDM, Anatrace), 0.2% (*w/v*) cholesteryl hemisuccinate (CHS, Anatrace), cocktail inhibitor and solubilized at 4 °C with gentle agitation for 2 hours. The extraction was centrifuged at 39,000 × g (Beckman) for 60 minutes to remove insoluble components, and the supernatant was incubated with anti-Flag affinity resin (GenScript Co., Ltd.) at 4 °C for 90 minutes. The resin was pooled and rinsed with 50 mL of buffer A containing 50 mM HEPES pH 7.5, 300 mM NaCl, 1 mM EDTA, 0.02% (*w/v*) lauryl-maltose-neopentyl-clycol (LMNG, Anatrace) and 0.002% (*w/v*) CHS, 30 mL of buffer B containing 50 mM HEPES pH 7.5, 300 mM NaCl, 1 mM EDTA, 0.01% (*w/v*) LMNG and 0.001% (*w/v*) CHS (Anatrace). The protein was eluted using a buffer containing 50 mM HEPES pH 7.5, 300 mM NaCl, 0.01% LMNG, 0.001% CHS, and 500 μg/mL synthesized FLAG peptide (GenScript Co., Ltd.). The eluted protein was further purified by SEC using a Superose 6 increase 10/300 column (GE Healthcare, USA) pre-equilibrated with running buffer containing 25 mM HEPES pH 7.5, 150 mM NaCl, 0.5 mM EDTA, 0.01% (*w/v*) LMNG and 0.001% (*w/v*) CHS. The purity of the proteins was confirmed by SDS-PAGE and Coomassie blue staining. Peak fractions containing the target proteins were pooled, concentrated to a final concentration of 3.5 mg/mL using Millipore 50-kDa cut-off filters (Millipore, USA), and subsequently flash-frozen in liquid nitrogen for cryo-EM analysis.

**Expression and purification of secreted SIDT1 and SIDT2 extracellular domain** The SIDT1^ECD^ and SIDT2^ECD^ proteins were expressed using FreeStyle 293-F cells (Invitrogen). The cells were cultured in OPM-293 CD05 medium (OPM Biosciences co., Ltd.) at 37 °C with 5% CO_2_ in a ZCZY-CS8 shaker (Shanghai Zhichu Instrument co., Ltd.) at 120 rpm. Transfection was performed when the cell density reached 2.5 × 10^6^ cells per mL by using expression plasmids and polyethylenimines (PEIs, Polysciences, USA). Approximately 1.5 mg of plasmids were premixed with 4.5 mg of PEIs in 50 mL of fresh medium for 25 minutes before application. Transfected cells were harvested 72 hours post-transfection and the 6 × His-tagged protein in the supernatant was purified using Ni-NTA (GE Healthcare, USA) affinity chromatography. The resin was washed with buffer A containing 50 mM HEPES pH 7.5, 300 mM NaCl, and 20 mM imidazole, and the target protein was eluted with buffer containing 50 mM HEPES pH 7.5, 300 mM NaCl and 250 mM imidazole. The purified protein was concentrated and subjected to gel-filtration chromatography using a Superdex 200 increase 10/300 column (GE Healthcare, USA), pre-equilibrated with gel-filtration buffer containing 25 mM HEPES pH 7.5, 200 mM NaCl.

### Single-particle Cryo-EM sample preparation and data collection

The purified SIDT1 sample (3 μL) was applied onto an H_2_/O_2_ glow-discharged, 300-mesh R0.6/1.0 holey carbon copper grid (preprocessed with 0.1% poly L-lysine hydrobromide after glow-discharging) (Quantifoil), or an R1.2/1.3 amorphous nickel titanium alloy (ANTA) film grid (Nanodim)^44^, respectively. The grid was blotted for 2.5 s with a blot force of -1 for the copper grid or 3 s with a blot force of 0 for the ANTA grid, both at 8 °C and 100% humidity. Finally, the grids were plunge-frozen in liquid ethane using a Vitrobot Mark IV (ThermoFisher Scientific). Cryo-EM datasets were collected using a 300 kV Titan Krios microscope (ThermoFisher Scientific) equipped with a K3 Summit detector (Gatan), or a K2 Summit detector with a GIF Quantum energy filter. The micrographs were automatically collected using SerialEM^45^ in super-resolution mode, with a nominal magnification of 29 k × (105 k × for the ANTA dataset). For the copper grid dataset, the exposure time was set to 2.4 s with a total accumulated dose of 60 electrons per Å^2^, resulting in a final pixel size of 0.82 Å, and a total of 9,115 micrographs were collected with a defocus range comprised between -1.2 and -2.0 μm. For the ANTA dataset, the exposure time was set to 11.02 s with a total accumulated dose of 60 electrons per Å^2^, which yields a final pixel size of 0.82 Å, and a total of 4,445 micrographs were collected with a defocus range comprised between -1.2 and -1.8 μm.

The data collection strategy for SIDT2 was consistent with that for SIDT1, resulting in 4,917 micrographs for the copper dataset and 5,547 (2,387 + 3,160) micrographs for the ANTA photographs, respectively.

The statistics of cryo-EM data collection are summarized in Table 1.

### Cryo-EM data processing

All dose-fractioned images were motion-corrected and dose-weighted by MotionCorr2 software^46^ and their contrast transfer functions (CTF) were estimated using Gctf^47^ in RELION^48^. The following particle picking, extraction, 3D classification and 3D refinements were carried out in RELION, while the initial 2D classification, the final non-uniform refinement and the local resolution estimation were performed using cryoSPARC^49^. For SIDT1, a total of ∼3,854 k particles were extracted for the subsequent 2D classification. After several rounds of 3D classification, a ∼4.1 Å map with clear TMDs and ECDs was generated. The remaining 249,851 particles were further classified, with 149,089 particles being re-extracted with a pixel size of 1.35 Å. These particles were polished with C2 symmetry, generating a density map with an overall resolution of 3.75 Å, which has been further refined to 3.33 Å using the non-uniform refinement and local refinement in cryoSPARC. Based on this, we create masks to realign and optimize the ECD and TMD regions, respectively. After non-uniform refinement and local refinement with C2 symmetry, we obtained a local optimization map for ECD with an overall resolution of 3.47 Å and a local optimization map for TMD with an overall resolution of 3.60 Å.

The data processing strategy for SIDT2 was consistent with that for SIDT1. A total of ∼3,164 k (∼1,694 k from the copper grid dataset and ∼1,470 k from the ANTA grid dataset) particles were extracted from a total of 10,464 micrographs for subsequent classifications. Similarly, we obtained a final map using 201,729 particles with an overall resolution of 3.17 Å, as well as two local-refined maps with an overall resolution of 3.27 Å for ECD and 3.33 Å for TMD, respectively.

The full cryo-EM data processing workflow is described in Supplementary Figs. 2 and 3.

### Model building and structure refinement

To build the SIDT1 dimer structure, an initial structure model for human SIDT1 predicted by AlphaFold^40, 50^ was placed and rigid-body fitted into the cryo-EM electron density maps using UCSF Chimera^51^. The ECD and TMD regions of SIDT1 were individually built based on two local-refined maps wherein these regions were better resolved. Bulky residues such as Phe, Trp, Arg, Lys and Tyr were used as references for sequence assignment. The manual and automated model building were iteratively performed using Coot 0.9.6^52^ and real-space refinement in Phenix 1.20^53^.

The model building and structure refinement strategies for SIDT2 were consistent with that for SIDT1.

The data validation and model refinement statistics are summarized in Table 1.

### Bimolecular fluorescence complementation

The GFP-based BiFC technique was used to investigate the SIDT1 and SIDT2 oligomerization in living cells. Fusion constructs were generated using fragments derived from GFP, including GFP fragment 1 (GN, amino acids 1-173) and fragment 2 (GC, amino acids 155-238)^54^. Full-length SIDT1 and SIDT2 cDNAs were inserted upstream of the GN and GC fragments, respectively, to generate recombinant proteins (SIDT1/2-GN and SIDT1/2-GC). These fusion constructs were connected by a Flag tag and a 10-amino acid linker encoding (GGGGS)_2_. The coding sequences of the recombinant proteins were then inserted into the *BamHI* and *EcoRI* sites of the pcDNA3.1(+) vector (Invitrogen), and all clones were verified through DNA sequencing.

Monolayers of HEK293T cells cultivated on coverslips were co-transfected with equimolar amounts of SIDT1/2-GN and SIDT1/2-GC plasmids using Lipofectamine 3000 (Invitrogen). After 24 hours of transfection, the culture media was removed, and the coverslips were washed with PBS. Subsequently, the cells were fixed with 4% paraformaldehyde at room temperature (RT) for 10 minutes, followed by a 10-minute incubation with 0.5% saponin to permeabilize the cells. The coverslips were then washed three times with PBS and stained with 2-(4-Amidinophenyl)-6-indolecarbamidine dihydrochloride (DAPI) for 10 minutes at RT. Images were acquired using a Zeiss LSM 980 scanning confocal microscope and processed with ZEN software (Carl Zeiss).

### Electrophoretic mobility shift assay

The interaction between SIDT1^ECD^, SIDT1^ECD-Dimer^, and SIDT2^ECD^ with various miRNA fragments was investigated using EMSA. A forward ssRNA oligonucleotide labeled with Carboxyfluorescein (FAM) at the 5’ end was synthesized by GenScript Co., Ltd. and annealed with an equimolar amount of the reverse strand. For the EMSA, increasing amounts of protein (1, 2, 3, 4, and 5 µM) were incubated with approximately 2.5 µM FAM-labeled miRNA or dsmiRNA at pH 5.5. EMSA reaction buffer containing 30 mM NaCl, 40 mM Tris (pH 5.5, adjusted with acetate), 2.5 mM EDTA, and 5% glycerol (*v*/*v*) or 30 mM NaCl, 25 mM Tris-HCl (pH 8.0), 192 mM Glycine, 5% glycerol (*v*/*v*) for a total volume of 20 µL at RT for 30 minutes. The resulting products were then separated on 5% native acrylamide gels (37.5:1 for acrylamide: bisacrylamide) in 1 × Tris-acetate-EDTA (TAE) running buffer (pH 5.5) or 1 × Tris-Glycine (TG) running buffer (pH 8.0)^55^ under an electric field of 10 V/cm for about 1 hour on ice. The gel was visualized and analyzed using the Tanon-5100 Fluorescent Imaging System (Tanon Science & Technology).

### Size exclusion chromatography

All proteins employed in biochemical assays were purified as previously described. We utilized SEC to assess SIDT1^ECD^ and SIDT2^ECD^ oligomerization. The SEC analyses were performed with a Superose 6 increase 10/300 column (GE Healthcare, USA) or a Superdex 200 increase 10/300 column (GE Healthcare, USA) in the buffer containing 25 mM MES pH 5.5, 100 mM NaCl. The peak fractions of targets were confirmed by SDS-PAGE followed by Coomassie blue staining. Data analysis was performed using GraphPad Prism 9 (GraphPad Software, San Diego, CA, USA).

### Sedimentation velocity analytical ultracentrifugation

We used SV-AUC to determine the sedimentation coefficients of SIDT1^ECD^, SIDT2^ECD^, SIDT1^ECD-Dimer^, SIDT1^ECD^-dsRNA, and SIDT1^ECD-Dimer^-dsRNA, and to calculate their molecular weights. The SV-AUC experiments were performed in a Beckman Coulter XL-I analytical ultracentrifuge (Beckman Coulter Inc., USA) using a two-channel centerpiece equipped with an An-50 Ti rotor (Beckman Coulter Inc., USA). Each sample was diluted to achieve a final volume of 400 μL (with an A280 nm absorption of approximately 1.0) in buffers with varying pH levels: a pH 3.5 buffer containing 25 mM sodium acetate and 100 mM NaCl, a pH 5.5 buffer containing 25 mM MES and 100 mM NaCl, and a pH 7.5 buffer containing 25 mM HEPES and 100 mM NaCl. Samples were loaded into 120-mm double-sector aluminum centerpieces and run at a rotor speed of 40,000 rpm under vacuum. Absorbance data were collected at 20 ℃, with simultaneous measurements at 280 nm and 260 nm for the apo-form and oligonucleotide-bound complex, respectively. SV-AUC data were globally analyzed using the SEDFIT program^56^ and fitted to a continuous c(s) distribution model to determine the sedimentation coefficients and molecular mass of each peak. All data were generated using GraphPad Prism 9 and Illustrator software, and c(s) and molecular weight were plotted.

### Microscale Thermophoresis

The MST experiments were conducted using a Monolith NT.115 instrument (NanoTemper Technologies). Purified SIDT1^ECD^ and SIDT1^ECD-Dimer^ were first exchanged into a labeling buffer containing 25 mM NaHCO_3_ pH 8.3, 100 mM NaCl, and 0.05% Tween-20. After buffer exchange, the proteins were labeled with a RED-NHS Labeling Kit (NanoTemper Technologies) according to the manufacturer’s instructions. For the MST measurements, the labeled proteins were dialyzed and exposed to various pH conditions using buffers: pH 3.5 (25 mM sodium acetate, 100 mM NaCl, and 0.05% Tween-20), pH 5.5 (25 mM MES, 100 mM NaCl, and 0.05% Tween-20), and pH 7.5 (25 mM HEPES, 100 mM NaCl, and 0.05% Tween-20). Throughout the experiments, the labeled protein concentration was maintained at 5 nM. 16-step serial dilutions of dsmiR168a or ssmiR168a were prepared in the same buffer to ensure consistent buffer conditions. MST measurements were performed at a constant temperature of 25 °C, with a 5-second LED on-time followed by a 30-second MST on-time. The LED and MST power settings were optimized for each experiment to achieve optimal signal-to-noise ratios and minimize aggregation or adsorption effects.

Data were collected and analyzed using the MO. Affinity Analysis software (NanoTemper Technologies). To ensure the accuracy and reliability of the results, MST experiments were performed in triplicate, and the data were averaged to determine the final *K_d_* values.

### DSF

The thermal stability analysis was conducted using the Tycho NT.6 (NanoTemper Technologies). SIDT1^ECD^ and SIDT2^ECD^ were diluted to a final concentration of 0.25 mg/mL at different pH values (pH 7.5, 5.5, and 3.5) and analyzed in triplicates using capillary tubes. Protein unfolding was monitored by measuring the intrinsic fluorescence at wavelengths of 350nm and 330 nm while gradually increasing the temperature from 35 to 95 °C at a rate of 30 K/minute. The obtained data was analyzed, smoothed, and the derivatives were calculated using the internal evaluation features of the Tycho instrument.

### Cross-Linking Assay

SIDT1^ECD^ was placed in an acidic buffer with 20 mM MES pH 7.5, 100 mM NaCl. Next, equimolar amounts of dsmiRNA168a and SIDT1^ECD^ were co-incubated at RT for 30 minutes. Following the incubation, glutaraldehyde crosslinking was carried out on proteins, with and without dsmiRNA168a. SIDT1^ECD^ was cross-linked in 20 mM MES pH 7.5, 100 mM NaCl, glutaraldehyde (GA) cross-linker (Macklin, Cat#C10366284) was resuspended to 10% (*v*/*v*) and the usage concentrations of glutaraldehyde (GA) include a gradient series: 0%, 0.05%, 0.1%, 0.2%, 0.5%. Cross-linking was allowed to proceed for 30 minutes at RT. Cross-linked samples were quenched in 0.1 M Tris-HCl pH 8.0, resolved by Western blot.

### SIDT1 and SIDT2 ceramidase activity assay

SIDT1 and SIDT2 enzymatic activity were evaluated through Liquid Chromatography-Mass Spectrometry/Mass Spectrometry (LC-MS/MS) analyses for sphingosine detection and quantification, using D-erythrosphingosine (d18:1) (Cat# 860490P, Avanti Polar Lipids) as the standard. Ceramidase activity was assessed by incubating 1.8 μM of purified SIDT1 and SIDT2 proteins with 40 μM ceramide (d18:1/18:0) (Cat# 860518P, Avanti Polar Lipids) at room temperature for 30 minutes, in a solution of 25 mM HEPES pH 7.5, 150 mM NaCl, 0.02% (*w/v*) DDM, and 0.001% (*w/v*) CHS. Methanol was added to quench reactions, achieving a final concentration of 30%. Ceramidase activity was then quantified by comparing peak areas with sphingosine standards. Lipids were extracted from reaction samples using the previously reported method14. The lipid extraction from samples proceeded by adding a dichloromethane/methanol/water (2.5:2.5:2 *v/v/v*) mixture and centrifuging. The collected organic phase was dried under nitrogen, redissolved in 30 µL of methanol, and stored at -20 °C until LC-MS analysis.

The LC-MS analysis was conducted using an AB SCIEX TripleTOF® 4600 System (SCIEX, Framingham, MA, USA). Sample separation was achieved on an absolute AQ-G8 column (particle size 3 μm, 2.1 × 100 mm) (Waters) maintained at a constant temperature of 40 °C. The mobile phases utilized were eluent A (containing 0.1% formic acid) and eluent B (comprised of acetonitrile). The gradient program was set as follows: 40% A and 60% B at 0 min, maintained at the same composition until 2 minutes, then changed to 70% B at 5 minutes, 95% B at 7 minutes, held at 95% B until 10 minutes, and returned to 40% A and 60% B at 10.1 minutes. The flow rate was maintained at 0.4 mL/min. The auto-sampler temperature was set at 8 °C, and the injection volume was 5 µL. The mass spectrometer was operated in positive electrospray ionization mode. The instrumental parameters were set as follows: Ion Source Gas1 (GS1) and Ion Source Gas2 (GS2) were both set at 55 psi, Ion Spray Voltage Floating (ISVF) was set at 5500 V, the source temperature was 600 °C, the declustering potential was 80 V, and the collision energy was 10 eV. Data was captured in full scan mode over a mass range of 100-1300 m/z with a scan time of 0.18 s. For tandem mass spectrometry (MS/MS) analysis, the 8 most intense precursor ions from each survey scan were selected for subsequent fragmentation and analysis. Data processing was performed using Analyst® TF 1.7 Software (SCIEX, Framingham, MA, USA). Analytes were quantified by comparing the peak areas to those of an internal standard. System calibration and analysis of quality control samples were carried out regularly to ensure data reliability and maintain system stability throughout the analysis. Quantification of the ceramidase activities of SIDT1 and SIDT2 was performed using d-erythrosphingosine (d18:1) as a standard through high-resolution LC-MS/MS analysis. LC-MS/MS analyses were performed using a Water ACQUITY HPLC system coupled with a QTRAP® 6500+ mass spectrometer (AB SCIEX, Framingham, MA, USA). Samples were injected onto a Water HSS T3 (2.1 × 100 mm, 1.8 μm) using an 8.5-minute linear gradient at a flow rate of 0.4 mL/min for the positive/negative polarity mode. The eluents were eluent A (0.1% Formic acid-water) and eluent B (0.1% Formic acid-acetonitrile). The solvent gradient was set as follows: 50% B, 1 minute; 50-95% B, 1-3 minutes; 95% B, 3-5 minutes; 95-50% B, 5-5.5 minutes; 50% B, 5.5-8.5 minutes. QTRAP® 6500 + mass spectrometer was operated in positive polarity mode with a Curtain Gas of 40 psi, Collision Gas of Medium, IonSpray Voltage of 5500 V, Temperature of 450 °C, Ion Source Gas of 1:55, Ion Source Gas of 2:55. The acquired data was controlled and processed using Analyst® TF 1.7 software (SCIEX, Framingham, MA, USA) for instrument control and Multiquant 3.03 software (SCIEX, Framingham, MA, USA) for MRM data processing, respectively.

### Statistics and reproducibility

Statistical analyses were performed in Prism 9 software (GraphPad) using a one-site-specific binding model. The MST assays were performed in triplicates. The experiments are reproducible. Data are shown as mean ± SD. The resolution estimations of cryo-EM density maps are based on the 0.143 Fourier Shell Correlation (FSC) criterion.

### Reporting summary

Further information on research design is available in the Nature Portfolio Reporting Summary linked to this article.

## Data availability

The source data for the graphs and charts in the figures is available as Supplementary Data 1. The coordinates and EM map files for the SIDT1-overall, SIDT2-ECD, SIDT1 TMD, SIDT2-overall, SIDT2-ECD and SIDT2-TMD have been deposited in the Protein Data Bank (PDB) and the EM Data Bank (EMDB) under accession number PDB-8K13, PDB-8K1B, PDB-8K1D, PDB-8K10, PDB-8K11 and PDB-8K12, and EMDB-36785, EMDB-36791, EMDB-36792, EMDB-36782, EMDB-36783 and EMDB-36784, respectively. For materials requests, please reach out to the corresponding authors.

## Acknowledgments

The cryo-EM studies were performed at the Center for Biological Imaging (CBI) (http://cbi.ibp.ac.cn), Institute of Biophysics, Chinese Academy of Sciences, and the Bio-Electron Microscopy Facility of ShanghaiTech University. We thank Dr. Qianqian Sun and Dr. Fengjiang Liu for their kind help with cryo-EM technical support. The MST studies were performed at the National Key Laboratory of Pharmaceutical Biotechnology at Nanjing University for technical assistance. We thank Tongyang Zhu for his help with MST data collection. The AUC studies were performed at Central Laboratory, Jiangsu Province Hospital of Chinese Medicine. We thank Biqing Chen for her help with AUC data collection. We thank Dr. Na Li at Shanghai Synchrotron Radiation Facility (SSRF) for her assistance in data collection and processing. We thank Dr. Haitao Yang for his valuable discussion and support.

This work was supported by grants from the National Key Research and Development Program of China (2018YFA0507103), the National Natural Science Foundation of China (grant No. 32270139 to X.J.), the Strategic Priority Research Program of the Chinese Academy of Sciences (XDB 37040102 to FS), the National Key Research and Development Program (2021YFA1301500 to FS), the National Natural Science Foundation of China (32071187 to YZ), the National Key Research and Development Program (2019YFA0904101 to YZ), the program for Innovative Talents and Entrepreneur in Jiangsu, and the Fundamental Research Funds for the Central Universities.

## Author Contributions

X.J. and C.Z. conceived the project. L.Z., T.Y., H.G. and X.J. supervised the project and designed all experiments. L.Z., T.Y., H.X., Y. Liu, Y.Y. and M.Z. cloned, expressed and purified SIDT1 and SIDT2 proteins. L.Z., T.Y. and H.X. performed all biochemical experiments. H.G., C.Q., Y. Lu and Y.G. collected and processed cryo-EM data. H.G. and C.Q. built and refined the structure models. X.J., F.S., Y.Z., H.G., and H.C.N. analyzed and discussed the data. All authors analyzed the final data and contributed to manuscript preparation. L.Z., T.Y., H.G., C.Q., H.C.N. and X.J. wrote the manuscript.

