## Supplementary Information for "Cryo-EM structures of human SID-1 transmembrane family proteins and implications for their low-pH-dependent RNA transport activity"

**This file includes:**

Supplementary Figures 1–11

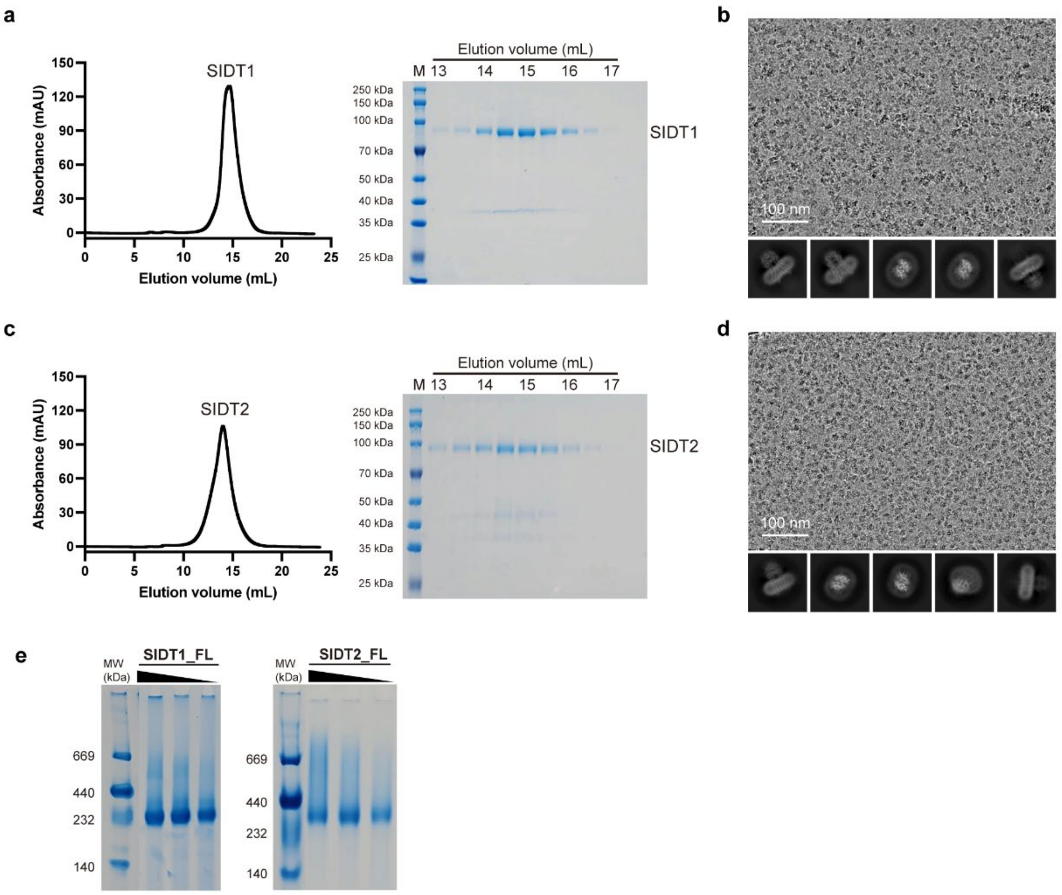


**Supplementary Fig. 1 | Protein purification and cryo-EM analysis of human SIDT1 and SIDT2. a, c** SEC profiles of SIDT1 (**a**) and SIDT2 (**c**) on a Superose 6 increase 10/300 column, respectively. Peak fractions were visualized by Coomassie blue staining. kDa, kilodaltons. M, marker. **b, d** Representative cryo-EM micrographs and 2D class averages of SIDT1 (**b**) and SIDT2 (**d**), respectively**.** The experiment was repeated independently more than three times with similar results. **e** BN-PAGE analysis of full-length SIDT1 and SIDT2, expressed in insect *Sf9* cells, reveals a homodimer formation in solution.

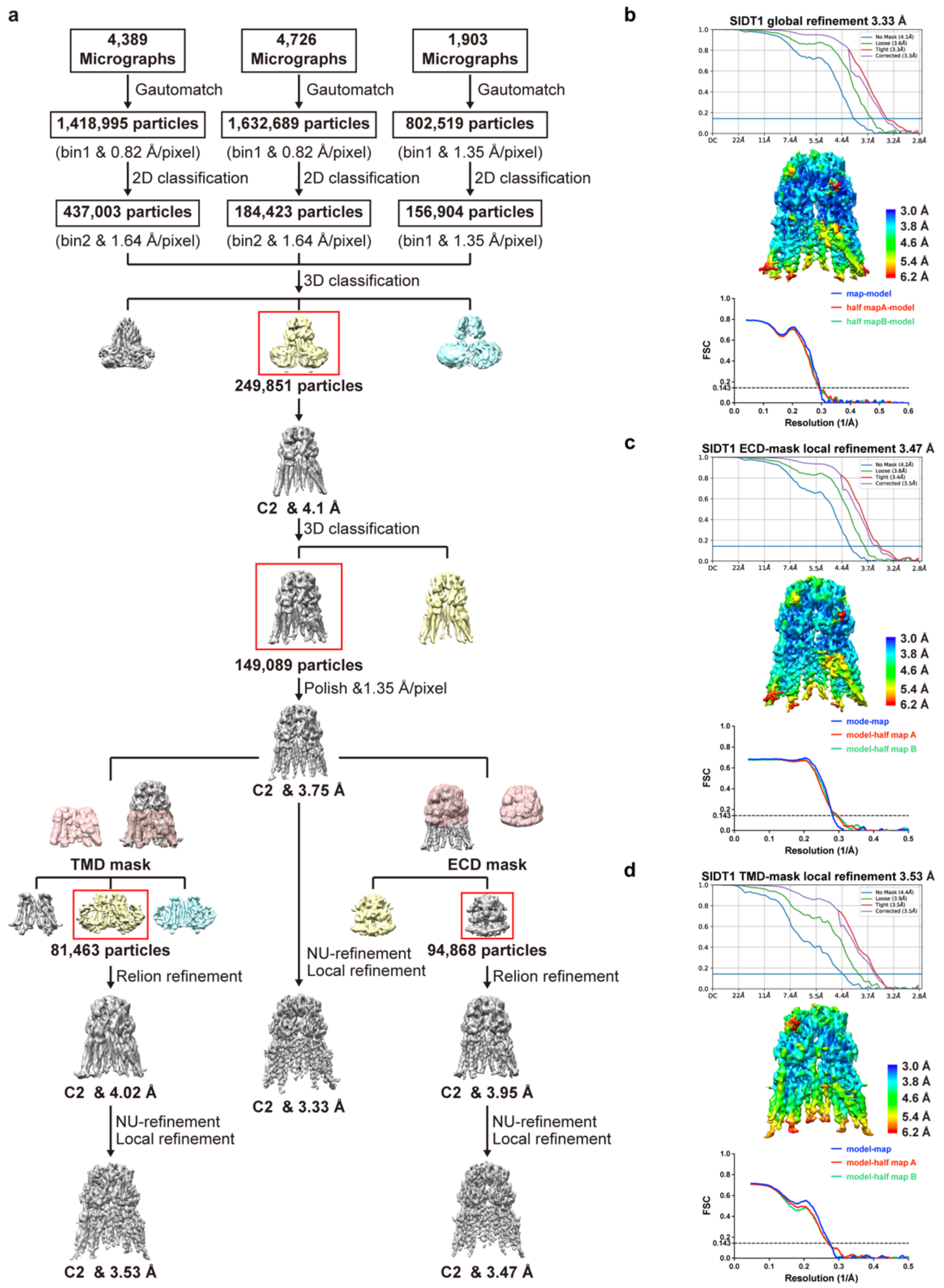


**Supplementary Fig. 2 | Cryo-EM data processing and validation for SIDT1. a** Workflow for the SIDT1 3D reconstructions. **b-d** The global FSC validation, cryo-EM density colored according to the local resolution and model-map FSC for the overall (**b**), local refinement with an ECD mask (**c**), and local refinement with a TMD mask (**d**) of SIDT1, respectively.

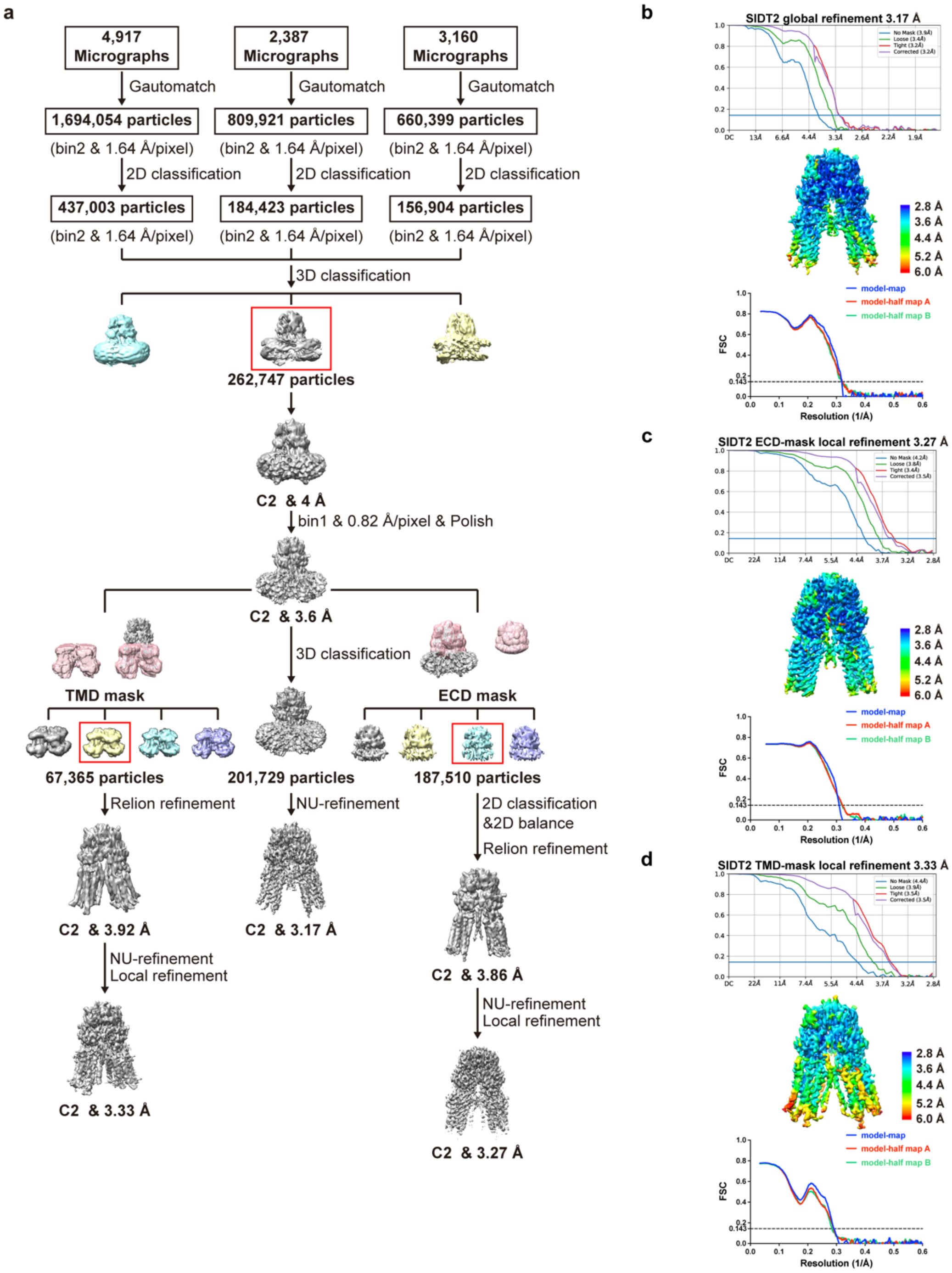


**Supplementary Fig. 3 | Cryo-EM data processing and validation for SIDT2. a** Workflow for the SIDT2 3D reconstructions. **b-d** The global FSC validation, cryo-EM density colored according to the local resolution and model-map FSC for the overall (**b**), local refinement with an ECD mask (**c**), and local refinement with a TMD mask (**d**) of SIDT2, respectively.

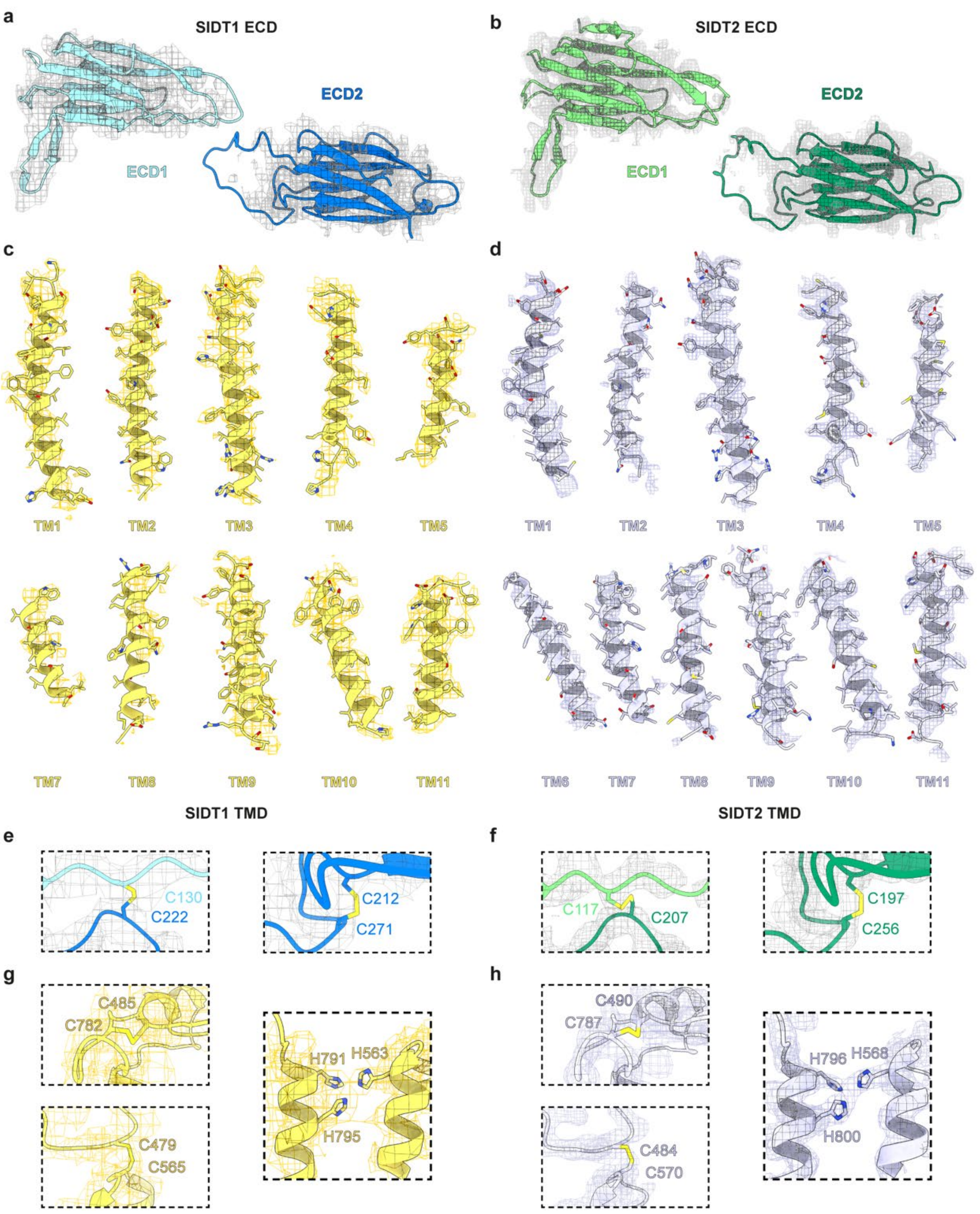


**Supplementary Fig. 4 | Representative sharpened cryo-EM density maps of SIDT1 and SIDT2. a, b** The EM densities of SIDT1 ECD (**a**) and SIDT2 ECD (**b**). **c, d** The EM densities of the TMD helices of SIDT1 (**c**) and SIDT2 (**d**). **e, f** The EM densities of two pairs of disulfide bond in SIDT1 ECD (**e**) and SIDT2 ECD (**f**). **g, h** The EM densities of two pairs of disulfide bond and the putative $Zn^{2+}$-binding site in SIDT1 TMD (**g**) and SIDT2 TMD (**h**).

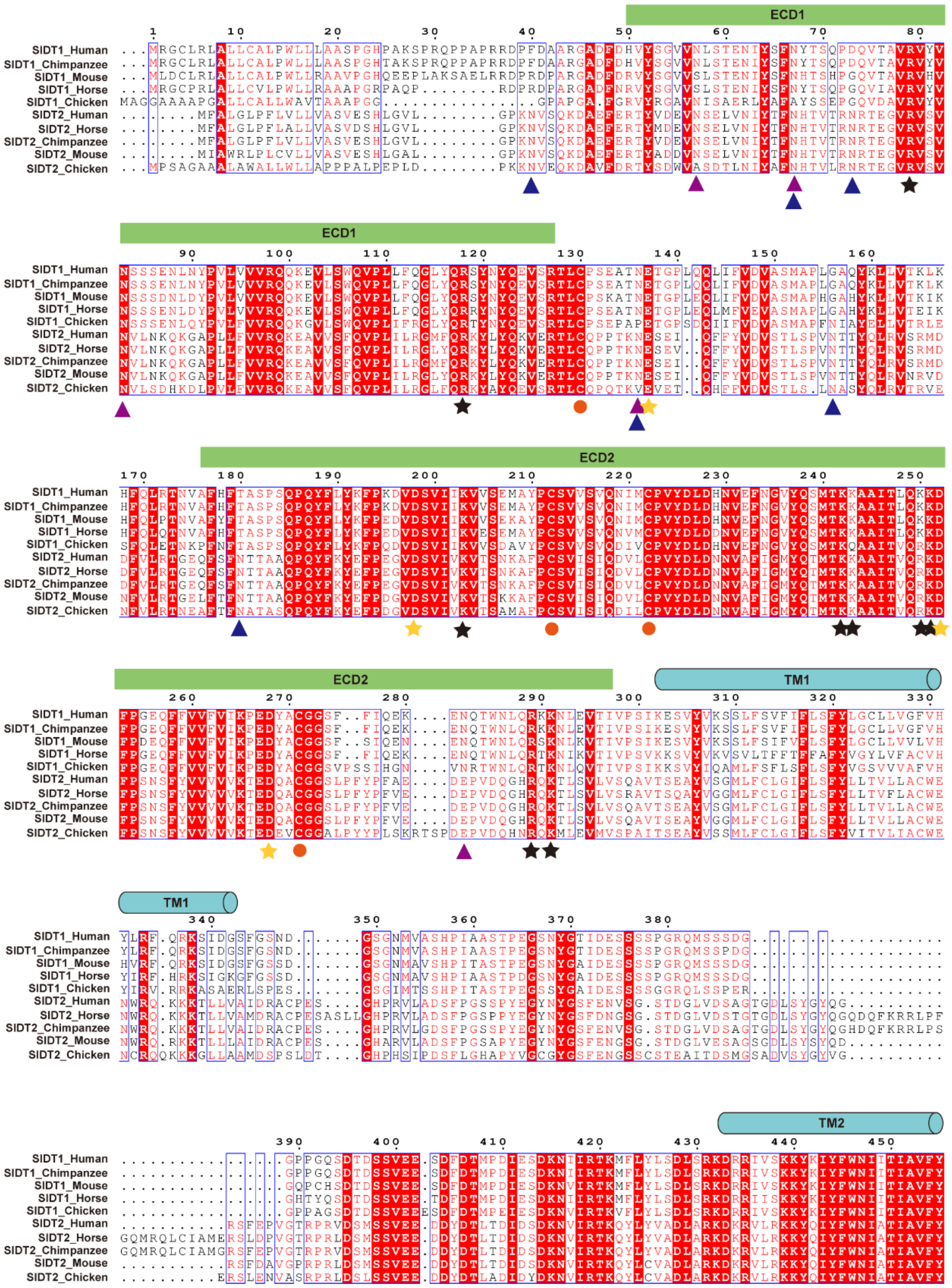
ECD1
ECD2
TM1
TM2
SIDT1_Human
SIDT1_Chimpanzee
SIDT1_Mouse
SIDT1_Horse
SIDT1_Chicken
SIDT2_Human
SIDT2_Horse
SIDT2_Chimpanzee
SIDT2_Mouse
SIDT2_Chicken

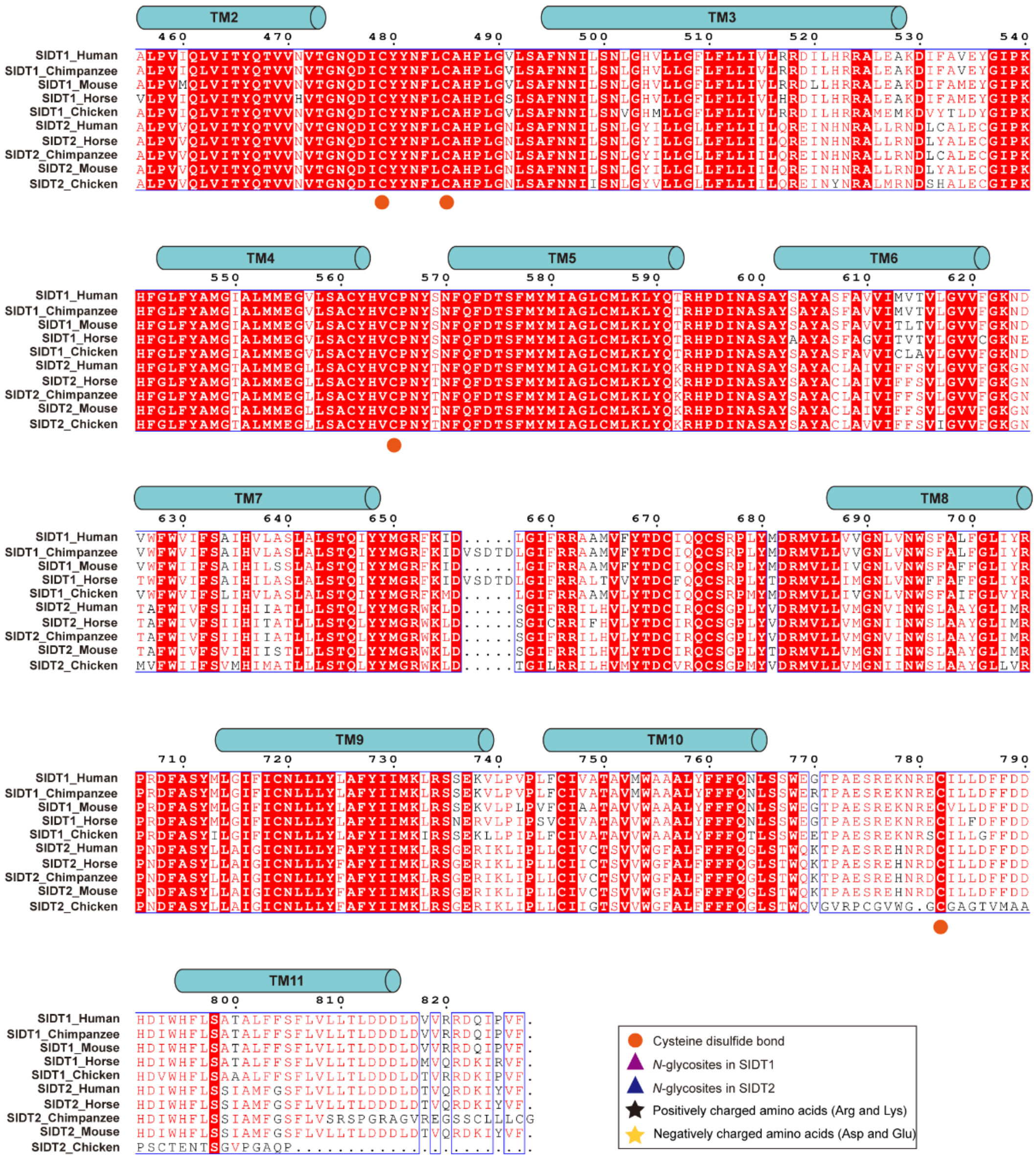


**Supplementary Fig. 5 | Sequence alignment of SIDT1 and SIDT2 with their homologs.** The primary sequences of different species are compared with that of SIDT1 using ClustalW[1] and visualized using ESPript3.0[2]. The secondary structural elements of SIDT1 predicted by AlphaFold[3] are indicated above the sequence alignment. Identically conserved residues are shaded red. Symbols below the alignment indicate cysteines for disulfide bond formation (orange circles), *N*-glycosites in SIDT1 (violet triangles), *N*-glycosites in SIDT2 (deep blue triangles), positively charged amino acids (blank rectangles), and negatively charged amino acids (yellow rectangles). The UniProt IDs for the aligned sequences are: SIDT1_chimpanzee: H2QN47;

SIDT1_mouse: Q6AXF6; SIDT1_horse: F6SIJ9; SIDT1_chicken: F1NBM4; SIDT2_chimpanzee: H2Q4U5; SIDT2_mouse: Q8CIF6; SIDT2_horse: F6Y6L2; SIDT2_chicken: A0A8V0ZU80.

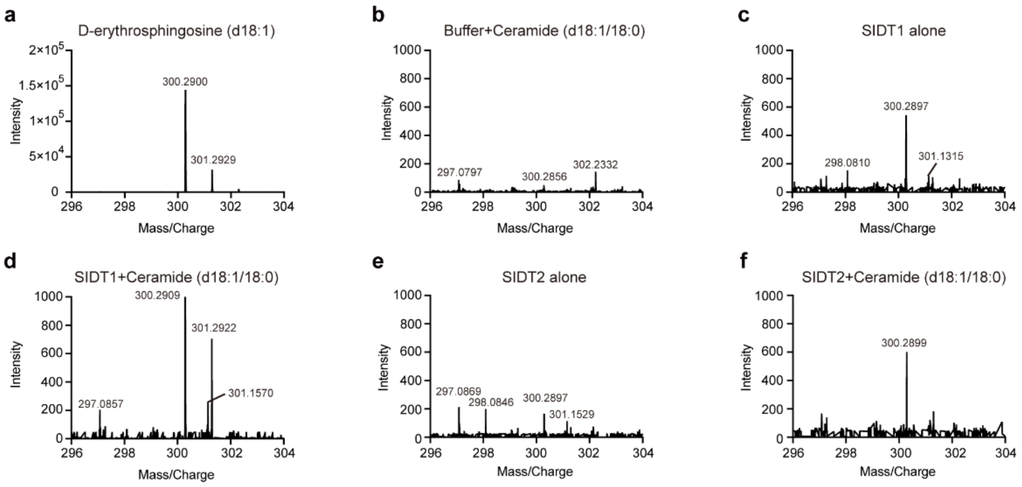


**Supplementary Fig. 6 | LC-MS/MS analysis of the ceramidase activity of SIDT1 and SIDT2**. **a-f** Representative mass spectrum for sphingosine detection is shown. The d-erythrosphingosine (d18:1) was used as the standard (**a**). Representative mass spectrum for the blank buffer plus ceramide (d18:1/18:0) (**b**), SIDT1 alone (**c**), SIDT1 plus ceramide (d18:1/18:0) (**d**), SIDT2 alone (**e**), and SIDT2 plus ceramide (d18:1/18:0) (**f**) are shown, respectively. The ceramidase activity of SIDT1 and SIDT2 was quantified by comparing the mass weight with a sphingosine standard.

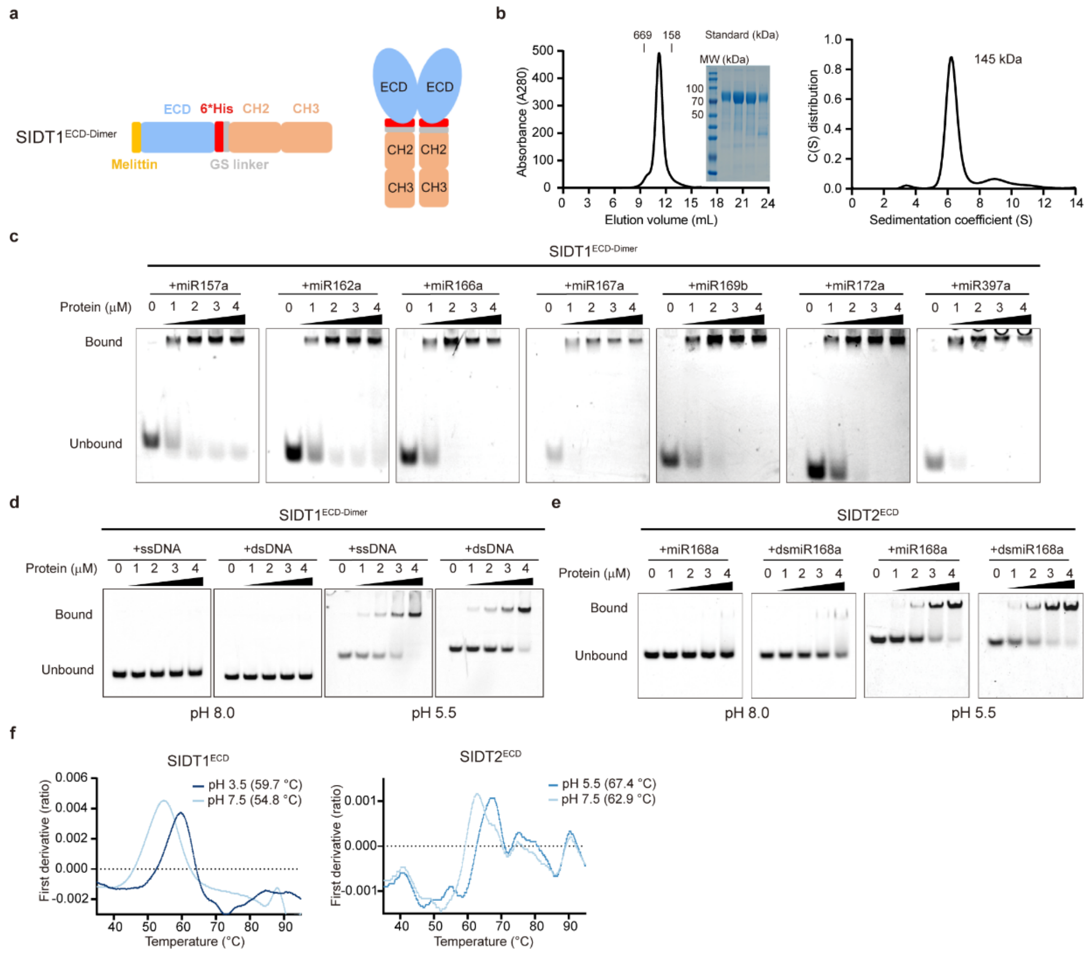


**Supplementary Fig. 7 | The ECDs of SIDT1 and SIDT2 bind to small RNAs under acidic conditions. a** Schematic diagram of a fusion protein of SIDT1$^{ECD}$ with immunoglobulin fragment (Fc; SIDT1$^{ECD\text{-}Dimer}$). **b** Characterization of the purified SIDT1$^{ECD\text{-}Dimer}$. Left panel: Chromatogram of purified SIDT1 $^{ECD\text{-}Dimer}$ via an Superose 6 Increase 10/300 column. SDS-PAGE gel (Coomassie stained) displays proteins in peak fractions. The calibration standard for gel filtration chromatography is a mixture of thyroglobulin, γ-globulin, and ovalbumin proteins with approximate molecular weights of 669 kDa (13.0 mL), 158 kDa (16.4 mL), and 44 kDa (17.5 mL), respectively. Right panel: SV-AUC analysis of the molecular weight of SIDT1$^{ECD\text{-}Dimer}$ in solution. This SV-AUC assay was performed twice with equivalent results. Values in the panel represent mean ± s.d. **c** An array of plant-derived miRNA-binding activities of SIDT1$^{ECD\text{-}Dimer}$ as revealed by EMSA at pH 5.5. The same EMSA setups as **Fig. 4c d, e** DNA binding activities of SIDT1$^{ECD\text{-}Dimer}$ (**d**), and miRNAs binding activities of SIDT2$^{ECD}$ (**e**) were examined in both pH 5.5 and pH 8.0 conditions. The same EMSA setups as **Fig. 4c** Bound, protein-miRNA complexes; Unbound, free miRNAs. **f** Protein stability of $SIDT1^{ECD}$ and $SIDT2^{ECD}$ was monitored in real-time as the temperature increased from 35 to 95 °C. DSF profiles (ratio between fluorescence at 350 nm and 330 nm) are displayed. DSF experiments were performed three times, and a representative result is shown.

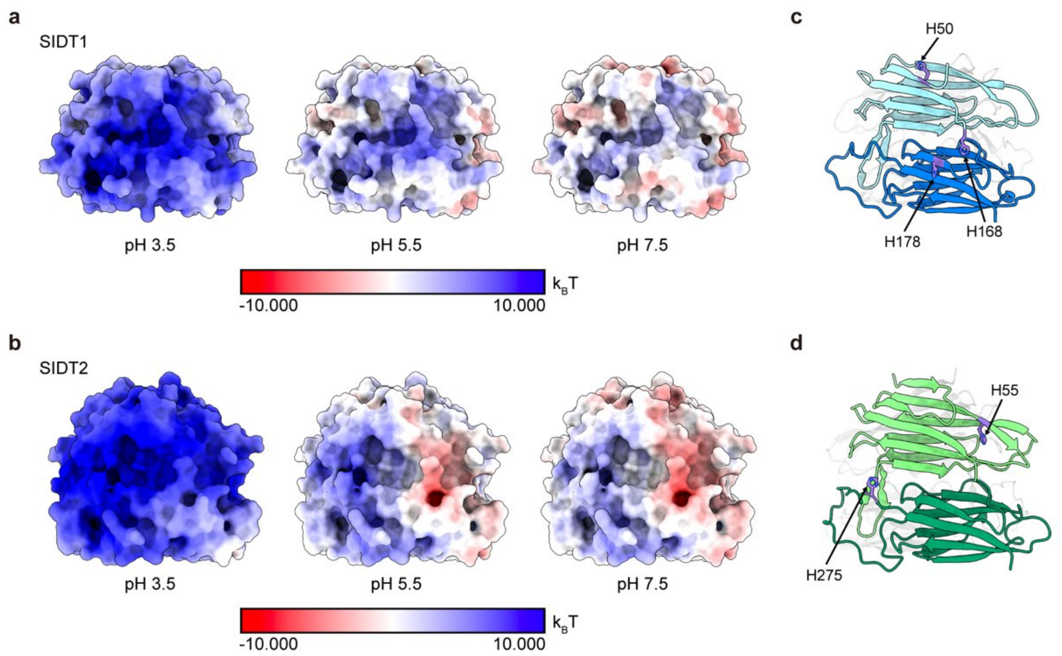


**Supplementary Fig. 8 | The protonation state of potential protein/RNA interface of SIDT1 and SIDT2. a, b** The electrostatic potentials of SIDT1 (**a**) and SIDT2 (**b**) at different pH values (pH 3.5, 5.5 and 7.5) are mapped onto their solvent-accessible surfaces. The electrostatic potentials are coloured from red (negative charge) to blue (positive charge) in the range of −10.0 to 10.0 $k_B T$. The electrostatic potentials of SIDT1 and SIDT2 were calculated using the APBS-PDB2PQR software suite[4] **c, d** The presence of histidine residues at the protein/RNA interface of SIDT1 (**c**) and SIDT2 (**d**) may contribute to the observed pH-dependent affinity. Histidine residues are shown as purple sticks.

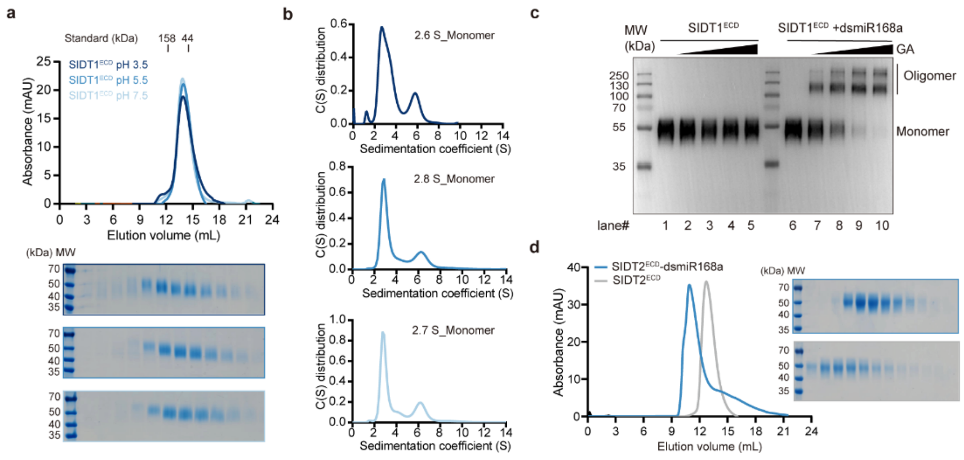


**Supplementary Fig. 9 | Small RNAs promote ECD oligomerization under acidic conditions. a** SEC and SDS-PAGE analysis of the oligomeric state of SIDT1$^{ECD}$ at pH 3.5 (deep blue), pH 5.5 (blue), and pH 7.5 (light blue), respectively. **b** SV-AUC analysis of the sedimentation coefficient corresponding to SEC results. SV-AUC experiments were performed twice with equivalent results. Values in the panel represent mean ± s.d. **c** Characterization of oligomeric state of SIDT1$^{ECD}$ via cross-linking assay. The cross-linking using glutaraldehyde (GA) and final products were analyzed by Western blot. Lanes 1 to 5 (without dsmiR168a) and 6-10 (with dsmiR168a) correspond to 30 minutes of incubation GA at pH 5.5 with at the following concentrations: 0%, 0.05%, 0.1%, 0.2%, and 0.5%. **d** The dsmiR168a triggers the assembly of SIDT2$^{ECD}$ dimer into oligomer, as determined by SEC.

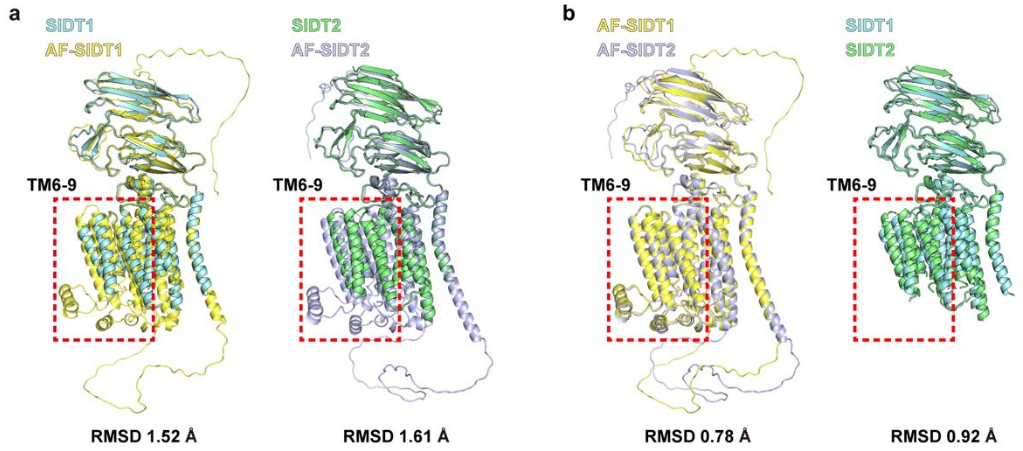


**Supplementary Fig. 10 | Structural comparison of the cryo-EM determined structures with the AlphaFold predicted structures. a** Comparison of experimental SIDT1 and SIDT2 models with their AlphaFold predicted AF-SIDT1 (left) and AF-SIDT2 (right) models, respectively. **b** Comparison between AlphaFold predicted models (left) and experimental models (right), respectively. RMSD, root-mean-square deviation.

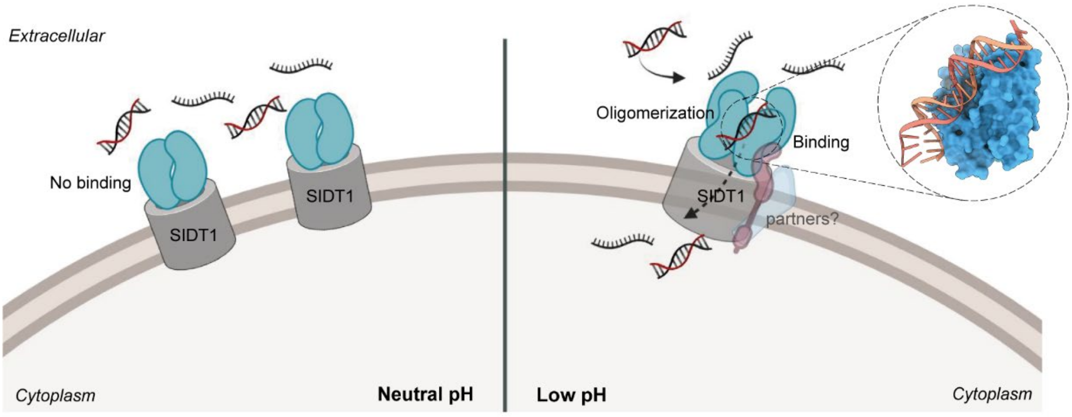


**Supplementary Fig. 11 | Possible working model depicting pH-dependent recognition by SIDT1 and their potential involvement in small RNAs transport.** This model is also applicable to SIDT2.

**Supplementary Table 1 | Sequences of plant-derived miRNAs prevalent in the serum of healthy Chinese individuals[5, 6].**

| Name | Sequence |
|---|---|
| miR157a | UUGACAGAAGAUAGAGAGCAC |
| miR162a | UCGAUAAACCUCUGCAUCCAG |
| miR164a | UGGAGAAGCAGGGCACGUGCA |
| miR166a | UCGGACCAGGCUUCAUUCCCC |
| miR167a | UGAAGCUGCCAGCAUGAUCUA |
| miR168a | UCGCUUGGUGCAGAUCGGGAC |
| miR169b | CAGCCAAGGAUGACUUGCCGG |
| miR172a | AGAAUCUUGAUGAUGCUGCAU |
| miR397a | UCAUUGAGUGCAGCGUUGAUG |
